# Aberrant DNA methylation distorts developmental trajectories in atypical teratoid/rhabdoid tumors

**DOI:** 10.1101/2022.03.14.483566

**Authors:** Meeri Pekkarinen, Kristiina Nordfors, Joonas Uusi-Mäkelä, Ville Kytölä, Minna Rauhala, Henna Urhonen, Laura Huhtala, Sergei Häyrynen, Ebrahim Afyounian, Olli Yli-Harja, Wei Zhang, Pauli Helen, Olli Lohi, Hannu Haapasalo, Joonas Haapasalo, Matti Nykter, Juha Kesseli, Kirsi J. Rautajoki

## Abstract

Atypical teratoid/rhabdoid tumors (AT/RTs) are pediatric brain tumors known for their aggressiveness, exceptionally low mutation rate, and aberrant but still unresolved epigenetic regulation. To evaluate methylation associated regulation in AT/RTs, we compared them to medulloblastomas and choroid plexus tumors by integrating DNA methylation (507 samples), gene expression (120 samples), and public transcription factor (TF) binding data. We showed that elevated DNA methylation masks the binding sites of TFs driving neural development and is associated with reduced transcription for specific neural regulators in AT/RTs. Part of the hypermethylated sites behaved similarly in AT/RTs and pluripotent stem cells, revealing DNA methylation -driven halted cell differentiation. AT/RT-unique DNA hypermethylation was associated with polycomb repressive complex 2 members, like EZH2, and linked to suppressed genes with a role in neural development and tumorigenesis. The obtained results highlight and characterize these DNA methylation programs as drivers of AT/RT malignancy.

## Introduction

Central nervous system (CNS) tumors can arise at any age and have the highest cancer-associated mortality rate in pediatric patients (1,2). Atypical teratoid/rhabdoid tumors (AT/RTs), medulloblastomas (MBs), and choroid plexus tumors (PLEXs) are CNS tumors detected in infants (3), though the latter two also have subtypes that occur in adults. AT/RTs and MBs are aggressive, grade IV tumors according to the World Health Organization (WHO) classification (4), and high-grade (III) PLEXs are typically detected in children (4–6). In particular, AT/RTs, non-WNT/non-SHH molecular group of MBs, and grade III PLEXs have a poor prognosis (7–9); thus, improving patient outcomes is an urgent task.

Because aberrant epigenetic regulation plays a major role in many aggressive pediatric tumors (10,11), we set out to specify the role of DNA methylation in AT/RT, and how it differs from MB and PLEX. For AT/RT, the significance of epigenetics is highlighted by their deserted DNA alteration landscape: the sole recurrent genetic alteration is the inactivation of *SMARCB1* or *SMARCA4*, both of which are subunits of mammalian SWItch/Sucrose-Non-Fermentable (SWI/SNF) chromatin remodeling complex (12). SWI/SNF complex is critical for the targeted opening of chromatin at certain genomic loci during neural development by recruiting acetyltransferase EP300, leading to histone 3 lysine 27 acetylation (H3K27ac) (12,13). Inactivation of SMARCB1 has been reported to activate stem cell-associated programs and to decrease the expression of neuronal differentiation genes in AT/RTs (11,14,15), thus promoting oncogenic gene regulation. In addition to AT/RT, SWI/SNF complex members are recurrently mutated in MBs (16,17). Polycomb repressive complex 2 (PRC2) is considered as an antagonist for SWI/SNF complex, as its key subunit EZH2 trimethylates H3K27 (leading to H3K27me3) and silences chromatin. However, H3K27me3 is also depleted in AT/RT (11), suggesting a H3K27me3-independent epigenetic driver for AT/RT development.

DNA methylation is one of the major mechanisms underlying epigenetic regulation. It contributes to cell differentiation (18); is altered in different malignancies (19–21); and is, together with histone methylation, a clinically relevant therapeutic target (22,23). In CNS tumors, DNA methylation is used for accurate tumor classification and early cancer detection (24–26). DNA methylation-based classification of CNS tumors also reveals tumor type-or subtype-specific DNA methylation patterns, linking DNA methylation to tumor type specific oncogenesis (26).

Traditionally, researchers have compared and analyzed subtypes of AT/RT, MB, and PLEX separately (27–30). For example, molecular AT/RT subtypes have been reported to have distinct genomic profiles, *SMARCB1* genotypes, and chromatin landscapes, including DNA methylation differences among the subtypes (27,31). Previous studies have also well characterized inter- and intra-tumoral heterogeneity in these malignancies. In this study, we set out to reveal hallmark DNA methylation patterns for AT/RTs and analyze their functional role in disease pathogenesis. MBs and PLEXs were used as external references and comparison points to facilitate tumor-type specific feature detection. AT/RT, MB, and PLEX also have a partly shared history and characteristics (32–34).

Our analysis revealed elevated DNA methylation patterns in AT/RT, suggesting that the binding of transcription factors (TFs) that drive neural differentiation is often inhibited in AT/RTs. In addition, PRC2 sites were hypermethylated in AT/RTs. We combined gene expression data with DNA methylation and observed upregulation of especially neural genes in MBs when compared with AT/RTs and/or PLEXs. Pluripotent stem cells (PSCs) and fetal brain (FB) were used as normal external references and to link DNA methylation changes to normal neural differentiation. We further analyzed two distinct DNA methylation programs in AT/RT, 1) PSC-like and 2) AT/RT-unique associated with PRC2 binding, to discover the key genes involved in DNA-methylation related oncogenic gene regulation.

## Results

### DNA is hypermethylated in AT/RT when compared with MB or PLEX

To study oncogenic epigenetic regulation in AT/RTs, we collected genome-wide DNA methylation Illumina microarray data (i450K) from altogether 497 AT/RT, MB, and PLEX tumors and expression data from 110 tumors; 89 normal brain DNA methylation samples were used as controls in i450k-based DNA methylation analysis. In addition, we generated data from 10 matched tumors with reduced representation bisulfite sequencing (RRBS) and RNA sequencing (**Figure 1A**) (**Supplementary Table 1**). DNA methylation data separated the tumor types and their subtypes in t-distributed stochastic neighbor embedding (tSNE) analysis (**Figure 1B**). MB samples were separated into WNT, SHH child/adult, SHH infant, group 3 and group 4 entities (26) based on DNA methylation and expression data. The tumor subgroups largely follow the diagnostic in the latest 2021 WHO classification (4), in which MB groups 3 and 4 form group non-WNT/non-SHH, MB SHH tumors are further divided based on TP53 mutation, AT/RTs are one entity, and PLEXs are divided into choroid plexus papilloma, atypical choroid plexus papilloma and choroid plexus carcinoma, which is also followed in DNA methylation -based subgrouping (4).

**Figure 1.**
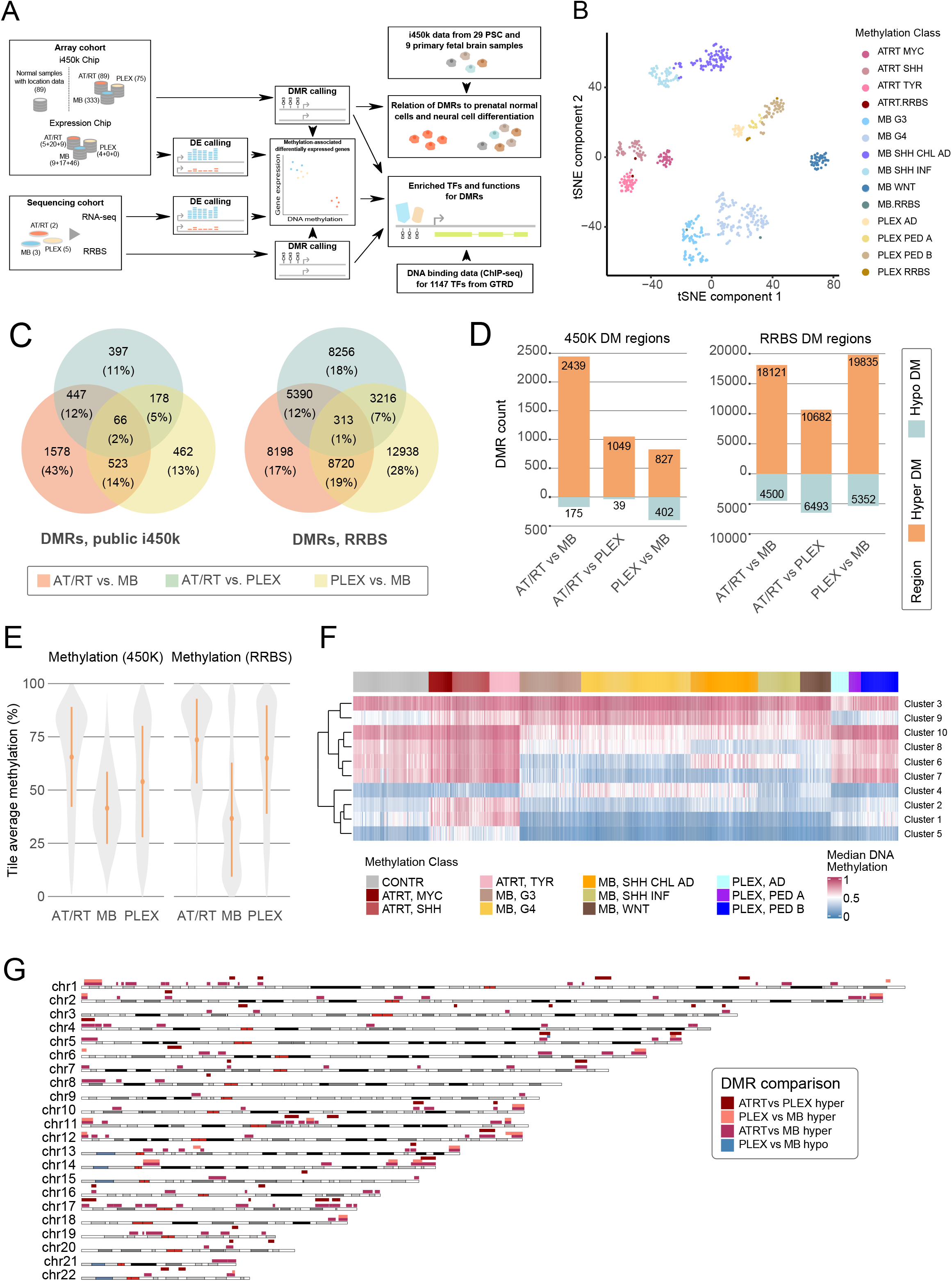
Characterization of DNA methylation differences among AT/RT, MB, and PLEX reveals AT/RT hypermethylation across all AT/RT subclasses and the genomic regions affected by large-scale DNA methylation changes. a) Illustration of our data analysis and integration approach. The number of samples in each cohort is shown on the left. Data was used to call differentially methylated regions (DMRs) and differentially expressed genes (DEs). Data from the gene transcription regulation database (GTRD) provided transcription factor DNA binding information. DNA methylation data from pluripotent stem cells and normal samples was used as references and to study normal neural cell differentiation. b) Tumor types are separated based on DNA methylation in a tSNE visualization, when the 10 000 most variable regions shared between i450k and RRBS data were used for the visualization. RRBS samples are positioned adjacent to the tumor subgroups that matched their clinical diagnosis. Methylation subgroups of each tumor type (omitted from (26)) in i450k were also visualized. c) Venn diagram showing the number of differentially methylated regions (DMRs) in each comparison. Tumor type-specific DMRs are marked into the intersecting areas. A higher number of DMRs were detected in the RRBS than in the i450k data. For i450k results, DMRs were filtered using methylation data from normal brain samples. d) For AT/RT, larger numbers of hypermethylated than hypomethylated regions were detected in all the comparisons both in i450k and RRBS data. Numbers of DMRs and direction of DNA methylation change for each comparison in both datasets. e) AT/RT had the highest overall DNA methylation level; MB was the least methylated. Violin plots visualized the distribution of DNA methylation in the 10 000 most variable sites in the RRBS and i450k microarray datasets. f) k-means clustering with 10 clusters revealed tumor type specific clusters. The median methylation of the regions in each cluster were used to summarize the methylation patterns. No AT/RT subtype specific clusters were detected but MB and PLEX both had clusters with differences between tumor subtypes. g) Several topologically associating domains (TADs) were influenced by large-scale methylation differences, especially in AT/RT-MB comparison. Karyoplot visualizes the TADs that harbor large-scale DNA methylation differences. Color indicates a comparison in which a difference was observed.

Differential methylation analyses resulted in 4,931 and 64,983 differentially methylated regions (DMRs) in the i450k (methylation difference >= 0.20, see details from Methods) and RRBS data (methylation difference >=0.25), respectively (**Figure 1C**). The RRBS DMRs were 1 kb segments and i450K DMRs had varying lengths, from 50-12349 bp. Higher DMR counts in RRBS data analysis were predictable because i450k DMRs were, on average, longer and RRBS has a wider genomic coverage (35,36). Furthermore, as explained in the Methods section, tumor location information and normal samples were used to filter out tumor location-related differences in i450k DNA methylation data, which decreased the DMR counts (**Figure 1C** vs. **Supplementary Figure 1A**). There was a higher proportion of intergenic DMRs in RRBS than in i450k data (36%, 30%, and 36% vs. 14%, 10% and, 11% in AT/RT-MB, ATRT-PLEX, and PLEX-MB, respectively) (**Supplementary Figure 1B, Supplementary Table 1**). DMRs were distributed across all chromosomes (**Supplementary Figure 1C**).

The majority of DMRs were more methylated in AT/RT than in other tumor types (84% and 72% in i450k and RRBS data, respectively) and less methylated in MB than in other tumor types (77% and 79% in i450k and RRBS data, respectively) (**Figure 1D**). Consistently, the highest DNA methylation levels were detected in AT/RTs and lowest in MBs (**Figure 1E**).

As higher DNA methylation levels were reported in AT/RT-TYR and AT/RT-SHH subgroups (31), we asked whether higher DNA methylation was subgroup-specific. No AT/RT subgroup specificity was observed in our setting; the highest DNA methylation levels were consistently detected in each AT/RT subgroup when compared with the subgroups of other tumor types (Wilcoxon Rank Sum p < 0.005, effect size Cliff’s delta |δ| > 0.2) (**Supplementary Figure 1D, Supplementary Table 1**). The k-means clustering of i450k DMRs resulted in 10 clusters (**Figure 1F and Supplementary Figure 2**), of which three were ATRT–specific (clusters 1, 2, and 5), one PLEX-specific (cluster 9) and two MB-specific (clusters 7 and 10). Clustering also revealed tumor subgroup -specific DNA methylation patterns, especially for MB (clusters 4, 6, and 8) and the AD subgroup of PLEX, but ATRT subgroups behaved similarly across clusters (**Figure 1F**) and also predominantly across individual methylation sites (**Supplementary Figure 2**).

DNA methylation can be focal or spread to larger genomic regions, and topologically associating domain (TAD) boundaries also serve as boundaries to the extension of epigenetic regulation (37). Indeed, our DMRs were involved in large-scale DNA methylation differences within specific TADs, especially in chromosomes 1, 11, 13, 17 and 19 in AT/RT-MB comparison (**Figure 1G**). Chromosome 17, which also harbors recurrent gains and losses in MB (38), carried 14 large-scale methylation-related TADs, of which 10 were differentially methylated in the AT/RT-MB comparison, 4 in AT/RT-PLEX comparison, and none in MB-PLEX comparison.

### Distinct neural differentiation related TFs are enriched to AT/RT and MB hypermethylated sites

To gather further information on the function and possible upstream regulation of cancer-specific DMRs, we used the chromatin immunoprecipitation sequencing (ChIP-seq) data collected from Gene Transcription Regulation Database (GTRD) (39), and aimed to find the TFs that had statistically significant over-representation of binding sites in cancer specific DMRs. We included only cancer-specific DMRs which behaved similarly in both comparisons (being *e*.*g*. hypermethylated in AT/RT when compared to both MB and to PLEX) and defined them as tumor-specific (**Figure 2A**).

**Figure 2.**
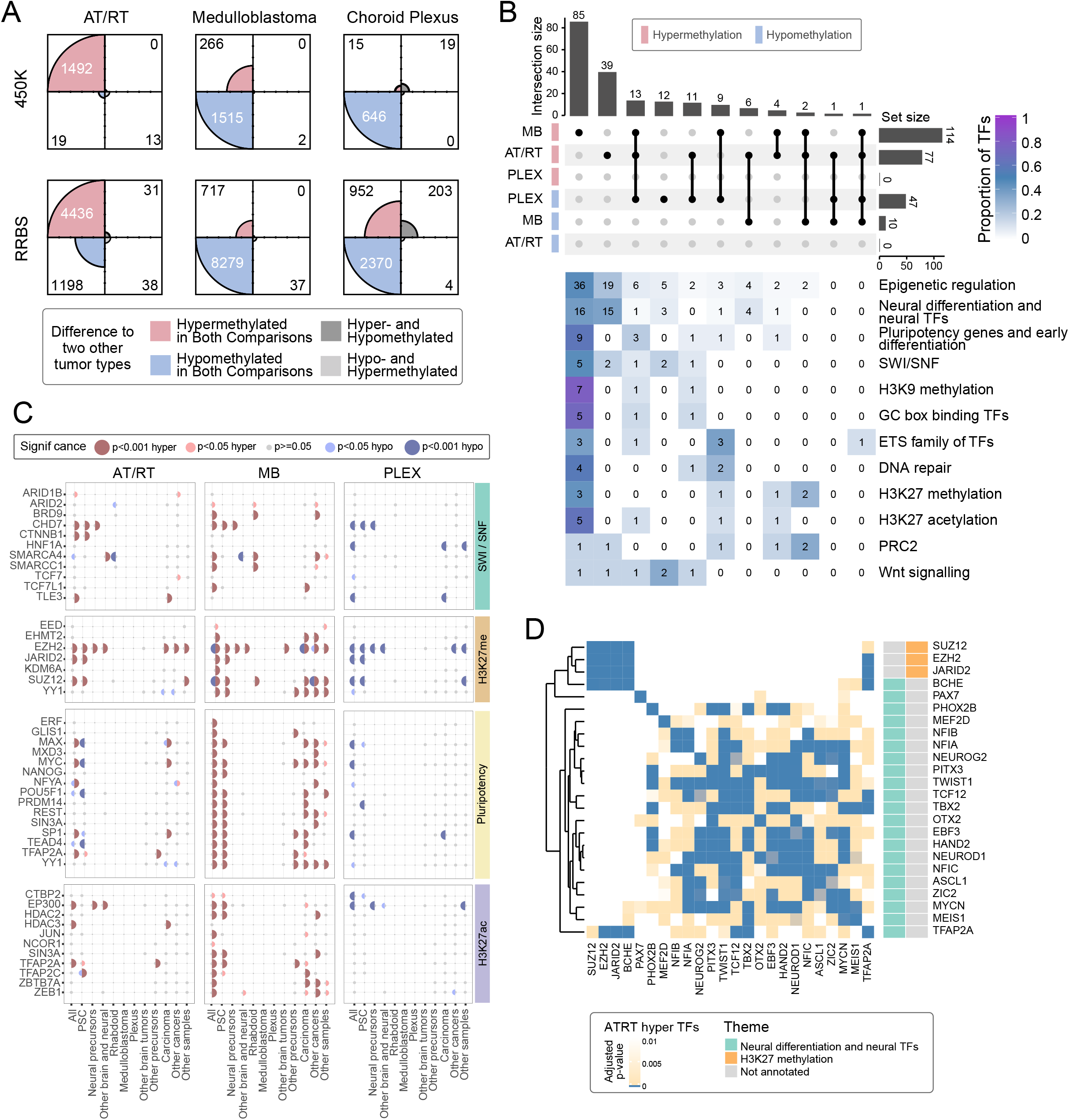
Neural cell differentiation -related TFs and key epigenetic regulators are enriched in tumor-type specific DMRs. a) The majority of AT/RT-specific DMRs were hypermethylated (99% and 79% in i450K and RRBS data, respectively), whereas hypomethylated DMRs were more commonly observed in MB (85% and 88% in i450K and RRBS data, respectively) and PLEX (98% and 71% in i450K and RRBS data, respectively). Four-field plots for each tumor type show the number of hypermethylated and hypomethylated regions in respect to two other tumor types. Tumor-specific DMRs have the methylation change into the same direction when compared with two other tumor types (*e*.*g*. hypermethylated in AT/RT when compared to MB and to PLEX). b) The majority of TFs are specifically enriched in AT/RT hyper, MB hyper, or PLEX hypo DMRs and largely linked to neural differentiation, SWI/SNF, and PRC2. Upper part: Upset plot of the number of enriched TFs for tumor-specific hypermethylated and hypomethylated regions. Some TFs were enriched to several DMR groups. Lower part: TFs were annotated into themes based on their reported functions. The number of TFs in a given theme for each upset plot column is shown. The color of the heatmap shows the fraction of TFs in each theme (row). c) TF binding in different sample types was variably associated with tumor-specific DNA methylation. GTRD data were split based on the sample of origin into the listed categories (at the bottom), and the association between TF binding and DNA methylation was calculated for each of the categories separately. TFs were organized into the themes listed in Figure 2B. The dot is not marked when a given TF is not measured in a given GTRD category. d) PRC2 subunits rarely co-localize with neural TFs in AT/RT hypermethylated sites. Heatmap visualization of the enrichment p-value (one-sided Fisher’s exact test) for colocalization. All the adjusted p-values of 0.01 or higher are marked in white. TF themes for each TF are annotated on the right-hand side of the heatmap.

Of the 183 enriched TFs, 136 (74%) were unique to one tumor type, 30 (16%) associated with two tumor types, and 17 (9.3%) to all three (**Figure 2B, Supplementary Table 2**). Altogether, 77 TFs were enriched in AT/RT hypermethylated DMRs (adjusted p-value < 0.05, one-sided Fisher’s exact test) and none in AT/RT hypomethylated sites (**Figure 2B**). For MBs, there were only 10 TFs enriched in hypomethylated DMRs and 114 TFs in hypermethylated DMRs, which is somewhat surprising as a moderate number of DMRs were hypermethylated in MB (**Figure 2A**). We hypothesized that these MB hypermethylated DMRs represent regulatory hubs that comprise several TF binding sites. Indeed, a higher proportion of MB hypermethylated DMRs was located in CpG islands than that for MB hypomethylated, AT/RT hypermethylated, or PLEX hypomethylated DMRs (**Supplementary Figure 3A, Supplementary Table 1**). No AT/RT subgroup specificity was observed in the DNA methylation patterns of differentially methylated TF binding sites (selected examples in **Supplementary Figure 3B**). DNA methylation reduces the DNA binding of most of the enriched TFs (23/36 TFs (64%) and 10/13 TFs (77%) with curated information in hyper- and hypomethylated regions, respectively) (40).

We used a literature search to functionally annotate the enriched TFs and used this information to divide 105/183 (57%) TFs into different themes (**Supplementary Table 2**). Themes were chosen based on the knowledge of these tumors, reported TF functions, and recurrent detection of certain gene family members (11,29,30,41). Notably, 15 of 39 (38%) TFs, which were uniquely enriched in AT/RT hypermethylated DMRs, were linked to neural differentiation and neural cells (**Figure 2B**). These included *e*.*g*. NEUROD1, ASCL1, and MYCN. When reanalyzing the TF binding after dividing the DMRs into clusters (**Figure 1F**), AT/RT-unique TFs BCHE, MYCN, NFIB, NFIC, and PITX3 were enriched to AT/RT-specific clusters 1 and/or 5 but none in AT/RT-specific cluster 2 (**Supplementary Table 2**). Four neural differentiation-related TFs (NEUROG2, HAND2, PHOX2B, and PAX7) were enriched in both AT/RT hypermethylated and MB hypomethylated DMRs (**Figure 2B, Supplementary Table 2**). Out of these, NEUROG2 and HAND2 were enriched only to cluster 2, and PHOX2B and PAX7 enriched to clusters 1 and/or 5. (**Figure 1F, Supplementary Table 2**).

We also noticed that 16 neural differentiation-related TFs (19% of the total 85) were uniquely enriched in sites hypermethylated in MBs (**Figure 2B**). However, five of these (REST, SIN3A, ZEB1, TBX1, and ZHX2) inhibit neural differentiation (42–45). In addition, four of the remaining 11 TFs were active in the later phases of neural differentiation namely during differentiation from neural precursors, whereas seven TFs hypermethylated in AT/RTs were active in earlier stages, namely during differentiation from pluripotent stem cells and during differentiation from neural stem cells (46–56). Finally, among the TFs uniquely enriched in MB hypermethylated regions, 9 TFs (11%) were related to the pluripotent stem cell state and early differentiation (**Figure 2B, Supplementary Table 2**) (42,57–59). Together these findings suggest that DNA methylation reduces the DNA binding of neural TFs, and thus neural differentiation, in AT/RT, whereas a more advanced neural differentiation is allowed in MB together with methylation-related silencing of pluripotency-related TF binding sites.

The expression of 164 (90%) enriched TFs was not significantly associated with any tumor type (**Figure 2B, Supplementary Figures 4 and 5, Supplementary Table 3**). ERF was the only differentially expressed (DE) TF in AT/RTs compared to other tumors (adjusted p < 0.01, absolute logFC > 1), whereas 13 TFs, including EBF3 and NEUROD1, showed MB-specific differential expression. Neural differentiation TFs enriched in AT/RT hypermethylated regions were expressed in AT/RTs and the highest expression of *e*.*g*. NEUROG2 and HAND2 were detected in AT/RTs (**Supplementary Figure 5A**). Thus, DNA methylation changes in TF binding sites affected the function of these TFs independently of their gene expression.

To analyze the TF binding data in more detail, we divided GTRD into 11 categories based on the cell of origin (**Figure 2C, Supplementary Figure 6, Supplementary Table 2**) (39). Using this split GTRD data, we found 15 neural differentiation and neural cells -related TFs measured in brain tumors or neural cells (**Supplementary Figure 6**). Out of these 15 TFs, 10 (67%) were enriched in AT/RT hypermethylated sites, MYCN (6.7%) in AT/RT hypomethylated sites, and only ZEB1 (6.7%) in MB hypermethylated regions (**Supplementary Table 2, Supplementary Figure 6**). On the other hand, neural TFs enriched only in MB hypermethylated sites were largely (12 of 16 TFs, 75%) measured from pluripotent cells or non-neural cell types (**Supplementary Table 2, Supplementary Figure 6**). Finally, TFs related to pluripotency and early cell differentiation were enriched especially in the MB hypermethylated regions (**Figure 2C, Supplementary Table 2**). All 11 pluripotent TFs, measured from pluripotent cells, were enriched in regions hypermethylated in MB. These together suggest that neural differentiation -related TF-binding sites in normal and other brain tumor cells are masked by DNA methylation in AT/RTs, whereas pluripotency-related TF binding sites are methylated in MB.

### DNA methylation is associated with SWI/SNF, PRC2 and acetyltransferase EP300 binding in a tumor-type-specific manner

As SWI/SNF chromatin remodeling complex is incomplete in AT/RTs due to SMARCB1 or SMARCA4 inactivation, we separately analyzed its subunits in GTRD data. Interestingly, SWI/SNF subunits behaved differentially: the binding sites of SMARCA4 and PBAF-specific ARID2 measured from rhabdoid tumor cells were enriched to hypomethylated AT/RT regions, but the binding sites of BRD9, the subunit of non-canonical complex lacking SMARCB1, were not associated with AT/RT-specific methylation (**Figure 2C**) (60). The binding sites of SMARCA4 in other brain and neural cells were enriched in sites hypermethylated in AT/RT and sites hypomethylated in MB (**Figure 2C, Supplementary Table 2**,). Consistently, the acetyltransferase EP300 binding sites in normal neural precursor and differentiated cells were hypermethylated in AT/RT, and EP300 binding sites in pluripotent cells were hypermethylated MB (**Figure 2C**). These results suggest that the neural SWI/SNF-EP300 target sites have been methylated in AT/RT. Furthermore, SWI/SNF binding sites are partly unique in MB and AT/RT, but the complex remains able to promote open chromatin in the regions to which it binds even in the absence of *SMARCB1*.

As PRC2 and EZH2 are antagonizing SWI/SNF and have been reported to populate neural SWI/SNF binding sites in AT/RT (11), we analyzed the association of PRC2 members with tumor-type-specific DMRs. Five of six (83%) TFs involved in PRC2 function and all seven (100%) TFs involved in H3K27 methylation (61–63) were enriched in sites hypermethylated in AT/RT or MB (**Figure 2B**). EZH2 was enriched in DMRs hypermethylated in AT/RT, hypermethylated in MB, or hypomethylated in MB, suggesting different modes of regulation (**Figure 2C, Supplementary Table 2**). When we analyzed TF binding in AT/RT hypermethylated regions after dividing them into DMR clusters (**Figure 1F**), EZH2 was enriched in cluster 1, which shows highest DNA methylation in AT/RT and lowest in MB (**Supplementary Table 2**). When examining split GTRD-data, all the EZH2 binding sites, but those originating from other brain tumors, were enriched in regions hypermethylated in AT/RT (**Figure 2C**). EZH2 was also enriched in regions hypermethylated in MB irrespective of the sample source.

### PRC2 subunits and neural differentiation -related TFs rarely co-localize

We performed TF binding co-localization analysis for regions hypermethylated in AT/RT or MB, or hypomethylated in MB or PLEX (**Figure 2D, Supplementary Figures 7 and 8**). Most TFs related to neural differentiation share their binding sites and are clustered, but significant co-localization (adjusted p < 0.01, one-sided Fisher’s exact test) was rarely observed between PRC2 members and TFs related to neural differentiation (**Figure 2D, Supplementary Figures 7 and 8**). In regions hypermethylated in AT/RT, the only TFs co-localizing with PRC2 subunits were the neural TFs BCHE and TFAP2A, which were co-localized with all the PRC2 subunits.

TFAP2A is known to regulate neural crest induction and cranial placode specification (64,65). BCHE is expressed early in nervous system development and plays a role in neural stem cell development (51). Only co-localization with one PRC2 subunit was detected for MBD1 and PHOX2B in MB hypermethylated and hypomethylated regions, respectively (**Figure 2D, Supplementary Figure 7**). The results suggest that PRC2-related DNA methylation largely affects parts of the genome other than the sites that recruit neural TFs during differentiation, BCHE and TFAP2A representing possible TFs that act at their intersection.

### DNA methylation regulates genes involved in neural cell differentiation

To associate DNA methylation with expression, we calculated DE genes (adjusted p < 0.01, minimum two-fold change) between tumor types in RNA-sequencing data and three public microarray datasets. Microarray expression datasets represented all the MB subgroups with similar subgroup proportions as in the DNA methylation microarray data (**Supplementary Tables 1 and 3**) (66). We used Venn diagrams to summarize genes that were similarly regulated in both microarray and sequencing experiments (**Figure 3A**). We next defined DM-DE genes i.e. the genes having an opposite direction in DNA methylation and gene expression change. Altogether 52 (9.4%), 41 (17%), and 56 (6.4%) DE genes harbor at least one DMR in the genomic neighborhood in the AT/RT-MB, AT/RT-PLEX, and MB-PLEX comparisons, respectively. There were 42 DM-DE genes with DNA methylation differences in the gene promoter and 67 in the gene-annotated enhancer region, respectively. The majority of the DM-DE genes (142 of 160 genes, 89%) were detected in only one comparison. In addition, a parallel increase/decrease in both expression and DNA-methylation was detected in 130 genes out of which 70 were in AT/RT comparisons (62 in AT/RT-MB and 8 in AT/RT-PLEX comparison, representing different genes) (**Supplementary Table 3**).

**Figure 3.**
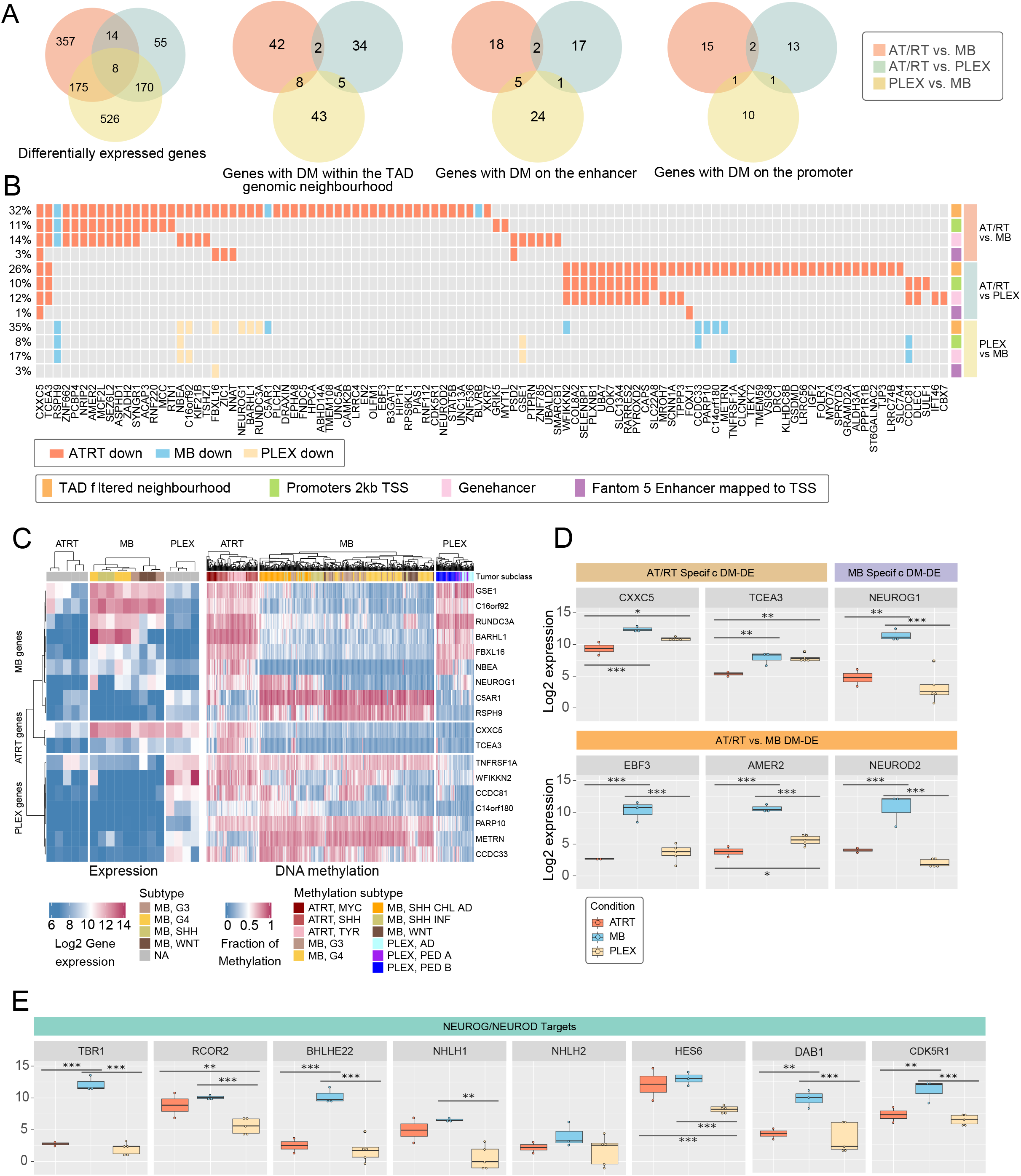
Methylation differences regulate both directly and indirectly genes relevant for neural differentiation and oncogenesis. a) Differential methylation (DM) was associated with differential expression (DE) of several genes. Gene expression and methylation patterns were studied in four contexts: 1) differential gene expression, 2) DE coupled with DM in the genomic neighborhood (+-200kb from the transcription start site [TSS] within the same topologically associating domain [TAD]), 3) DE coupled with DM in gene-linked enhancer, and 4) DE coupled with DM in the gene promoter (2kb upstream and 500bp downstream from the TSS). Venn diagrams show the genes that behaved similarly in both sequencing and array data. Differentially expressed genes associated with differential methylation (cases 2-4) are called DM-DE genes. Only cases where the sign of DM change was opposite to DE were included in the figure. b) The DM-DE genes were hypermethylated and underexpressed in AT/RT. Oncoprint showing the DM-DE genes, which are differentially expressed in AT/RT-PLEX and/or AT/RT-MB comparisons, and the direction of differential gene expression. For each comparison, different possible DMR locations (gene neighborhood within TAD, promoter (2kb), enhancer (Fantom 5), enhancer (Genehancer)) are visualized separately on the right side of the figure. Enhancer regions have been linked to the DE genes. c) Sample-wise levels of gene expression (on the left) and DNA methylation (average methylation of variable sites, right) for tumor-specific DM-DE genes in the microarray datasets. d) Expression patterns of selected DM-DE genes. \**p* < 0.05, \*\**p* < 0.01, *** *p* < 0.001 e) Expression of downstream targets of NEUROG and NEUROD TFs. The highest expression is typically detected in MB. \*\**p* < 0.01, *** *p* < 0.001

DM-DE genes were summarized using oncoprint-like visualization (**Figure 3B**, full version in **Supplementary Figure 9**). Nearly all DM-DE genes in AT/RT comparisons (101 of 104, 97%) were downregulated in AT/RT. In MB and PLEX, DNA methylation differences were more linked to increased expression: of all the 160 DM-DE genes, 80 (65% of MB-related), and 81 (73% of PLEX-related) were upregulated in MB and PLEX, respectively, when compared with at least one tumor type (**Supplementary Figure 9**). DM-DE genes were predominantly hypermethylated in AT/RT (141/160 genes (88%)) and overexpressed in PLEX or MB or in both in respect to AT/RT (120 and 112 genes in array and RNA-seq data, respectively) (**Supplementary Figure 10**). The number of AT/RT-specific genes was partly reduced because decreased DNA methylation in both MB and PLEX led to increased expression in only one of them.

Altogether 18 DM-DE genes were specific to a given tumor type, that is, differential methylation in the same genomic context (genomic neighborhood, promoter or enhancer) was observed in two tumor comparisons in both microarray and sequencing data (**Figure 3B-C, Supplementary Figure 9-10**). Of the four tumor type-specific genes that were hypermethylated and underexpressed, *CXXC5* and *TCEA3* were specific to AT/RT, and *RSPH9* and *C5AR1* specific to MB. Of the 14 tumor type-specific genes that were hypomethylated and overexpressed, seven genes, including neural genes *NEUROG1, NBEA* and *BARHL1*, were specific to MB. Of these, *BARHL1* has been previously reported to be overexpressed in MB (67).

We identified *NEUROG1* and *NEUROD2* as DM-DE genes that were hypermethylated in AT/RT (**Figure 3B-D**), and *NEUROG2* and *NEUROD1* as TFs whose binding sites were enriched specifically in AT/RT hypermethylated regions (**Figure 2D, Supplementary Figure 6**). Because these TFs are key regulators of neural differentiation (68,69), we visualized their expression together with the expression of their target genes *TBR1, RCOR2, BLHE22, NHLH1, NHLH2, HES6, DAB1*, and *CDK5R1* (**Figure 3E, Supplementary Figure 11**). Except for NEUROG2, which had the highest expression in AT/RT, the highest expression of all these genes was observed in MB. This result suggests that their induction, which should occur during neural differentiation, has been successful mainly in MB.

### AT/RTs harbor both unique and pluripotent stem cell -like DNA methylation patterns

DNA methylation is an important regulator of cell differentiation and our findings suggest that AT/RTs, MBs and PLEXs represent different developmental stages. As DNA methylation in AT/RTs has also been recently reported to show similarities to PSCs (70), we analyzed the observed DNA methylation differences in the context of normal neural cell differentiation by utilizing i450k DNA methylation data from PSC and primary FB samples (71,72) (**Figure 4A**). We defined whether the observed tumor type-specific DMRs in MB and AT/RT are unique to tumor type, PSC-like, or FB-like, and whether their DNA methylation level is changed during normal neural differentiation (**Figure 4B**, left side, **Supplementary Table 4**). Out of the 898 DMRs hypermethylated in AT/RT, 322 (39%) were similarly methylated in PSCs as in AT/RTs and hypomethylated during differentiation from PSCs to FB (group 6, **Figure 4B**). Only 31 (3.5 %) of AT/RT hypermethylated DMRs were FB-like (group 8), highlighting the higher similarity of AT/RTs to PSCs than FB. However, there were also 448 (50%) AT/RT-unique methylation patterns. Out of these, 124 (14%) were most methylated in AT/RT and not related to differentiation (group 4), 176 (20%) were most methylated in AT/RT and also hypermethylated in PSCs when compared to FB (group 1), and finally 148 (16%) of which were differentiation-related (with higher methylation in PSCs than FB) but also hypomethylated in AT/RTs when compared to PSCs (group 7) (**Figure 4B**).

**Figure 4.**
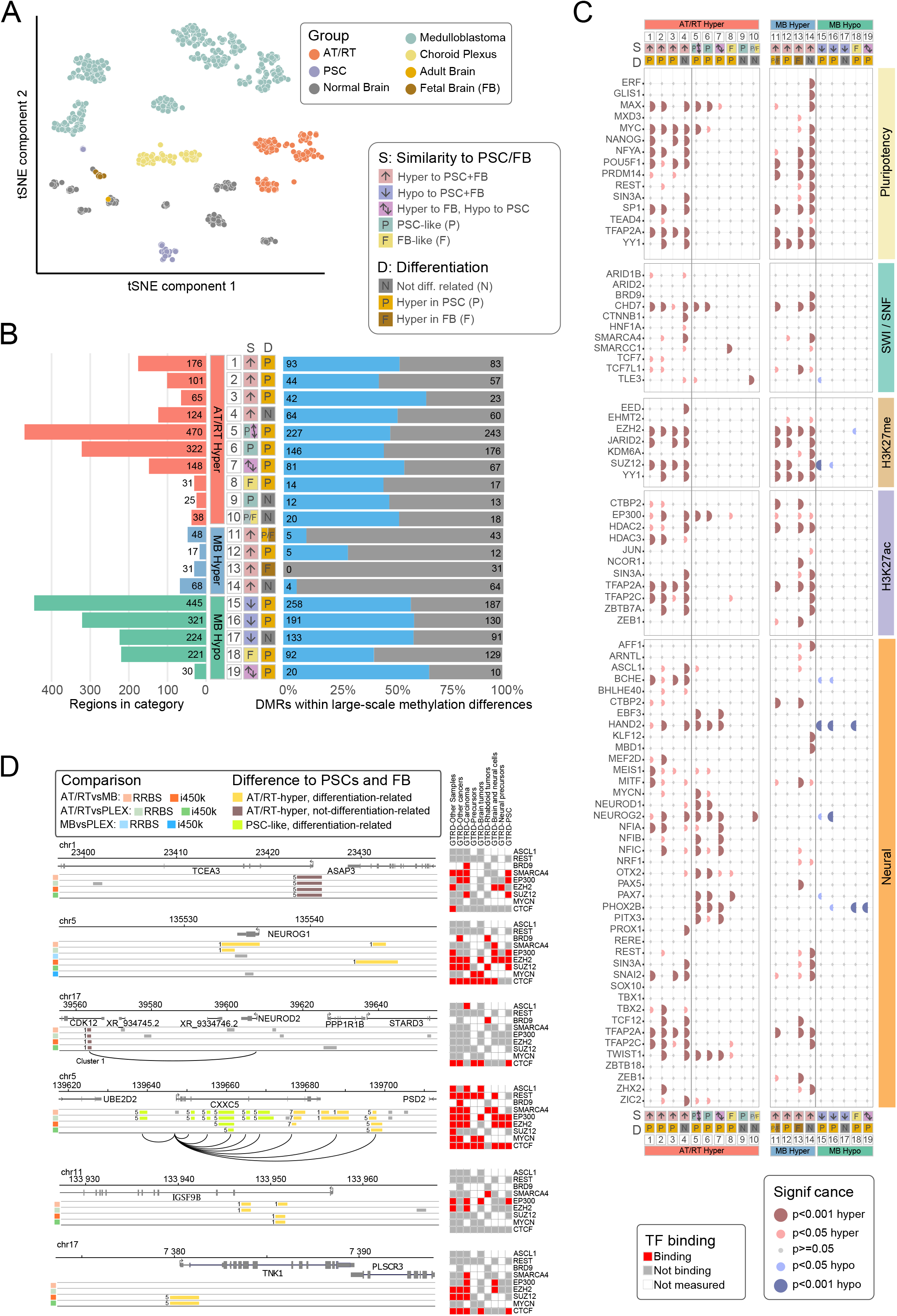
AT/RTs harbor pluripotent stem-cell-like and AT/RT-unique DMRs, which are associated with the suppression of relevant regulators. a) Pluripotent stem cells (PSCs), primary adult brain, and primary fetal brain (FB) are separated from tumor samples based on DNA methylation in a tSNE visualization, when the 10 000 most variable regions in i450k data were used for visualization. b) AT/RT hyper DMRs were mostly AT/RT-unique or PSC-like, whereas MB DMRs were MB-unique or FB-like. Very few MB hypermethylated DMRs were associated with large-scale differences in DNA methylation. Tumor-type-specific DMRs (Figure 2A) were categorized based on DNA methylation levels in PSC and FB samples. The bar plot on the left shows the number of DMRs in different categories. Annotations show whether DMRs are PSC-like (P), FB-like (F), or unique (different from PSC and FB) and whether or not they were differentially methylated between PSC and FB. AT/RT-unique DNA methylation groups 2 and 3 are subgroups of 1. Group 2 includes only DMRs which were also hypermethylated in PSCs compared to PLEX or MB. Group 3 includes AT/RT-unique DMRs that were also PSC-like in PLEX. Group 5 comprises groups 6 (PSC-like DMRs) and 7 (AT/RT-specific DMRs which are hypermethylated in PSCs compared to AT/RT). MB-unique hypermethylation group 11 comprises groups 12 (with higher methylation in PSCs than FB) and 13 (with higher methylation in FB than PSCs). Group 16, comprising MB-unique DMRs that are also PSC-like in AT/RT, is a subgroup of MB-unique group 15. The proportion of DMRs in large-scale methylation differences within annotated DMR categories is shown in blue on the right. The number of DMRs is marked in the figure. c) The DMR category-related binding patterns revealed TFs involved in tumor-unique, normal cell -like, and differentiation-related regulation of DNA methylation. PRC2 subunits were enriched to the AT/RT-unique DMRs wereas neural TFs were enriched to both AT/RT-unique and PSC-like DMRs with varying enrichment patterns. TF binding site enrichment was calculated separately for each DMR category (bottom). TFs were organized into the themes listed in Figure 2B. The dot is not marked when a given TF is not measured in a given GTRD category. d) Hypermethylated DMRs in relevant genes, which are suppressed in AT/RT. Visualization of DMRs around *NEUROG1, CXXC5, TCEA3, NEUROD2, TNK1*, and *IGSF9B. CXXC5* and *TCEA3* are AT/RT-specific suppressed DM-DE genes, *NEUROG1* and *NEUROD2* are DM-DE genes in AT/RT-MB comparison. *TNK1* and *IGSF9B* genes are connected to AT/RT-unique DMRs harboring EZH2 binding sites. Distal DMRs are connected to the TSS via an arch. Oncoprint indicates which relevant TFs have binding sites in these regions in selected GTRD categories. Color of the DMR indicates if the DMR is PSC-like and whether it is differentiation related. Number in front of DMR indicates the k-means cluster which DMR belongs to (see Figure 1F). Gray DMRs were not included in TF binding and DMR cluster analysis.

By contrast, 116 of 123 (94%) MB hypermethylated DMRs and 699 of 957 (73%) MB hypomethylated DMRs were unique to MB (**Figure 4B**). Furthermore, 241 (25%) MB hypomethylated DMRs were FB-like and hypomethylated during differentiation (group 18). Within MB specific regions, 445 of the 957 MB hypomethylated regions (46%) were hypomethylated during differentiation (group 15), 321 (34%) of which were also PSC-like when compared to AT/RT (group 16). A similar trend was observed for 123 MB-unique hypermethylated regions, of which 68 (55%) were not differentiation-related and 31 (25%) were hypermethylated during differentiation (group 13) (**Figure 4B**). Thus, a high proportion of even tumor-type-unique DMRs change their DNA methylation during differentiation and are differentiation-related in a manner that underlines the similarity of MBs to FB and AT/RTs to PSCs.

These results further support our prior findings that a majority of AT/RT-specific DMRs are hypermethylated in AT/RT when compared with MB and PLEX (**Figure 2A**). The majority (75%) of these sites are demethylated upon neural differentiation, and a large proportion (39%) of them are PSC-like (**Figure 4B**). However, we also observed AT/RT-unique DNA methylation, which represents unique targeting of DNA methylation in these malignant cells (**Figure 4B**).

We calculated the proportion of large-scale DMRs (**Figure 1G**) in each DMR category (**Figure 4B**, right side) to determine whether the observed DNA methylation differences were more linked to focal or large-scale regulation of DNA methylation. Notably only 14 (11%) of MB hyper DMRs represented large-scale DNA methylation differences, whereas 384 (49%) and 505 (55%) of AT/RT hyper and MB hypo DMRs, respectively, were involved in large-scale methylation (**Figure 4B**). Thus, MB hyper DMRs appear to target specific cytosines or focal areas, such as specific promoters and enhancers, whereas both focal and large-scale regulation of DNA methylation is observed for regions hypermethylated in AT/RT or hypomethylated in MB.

### AT/RT-specific methylation is associated with PRC2

We analyzed TF binding separately in the DMR categories presented in **Figure 4B** to define TFs relevant to different regulatory programs. The binding of EZH2 and other PRC2 members was significantly enriched in AT/RT-specific, but not in PSC-like or FB-like, AT/RT hyper DMRs (**Figure 4C**). In MB, PRC2 members were enriched in hypermethylated regions, but EZH2 was also enriched in hypomethylated regions that were FB-like. Interestingly, many of the AT/RT-unique differentiation-related DMRs with EZH2 binding site were linked to genes which are involved in dendrite growth and differentiation from neural cells to mature state, which are the key processes the neuronal BAF complex (nBAF, mammalian neuronal SWI/SNF complex) is activating during neural development (**Supplementary Table 4**) (73). We saw *e*.*g. IGSF9B* that is related to synapsis development (74). Two DM-DE genes, *DOK7* (upregulated in PLEX when compared to ATRT and MB) and *CAMK2B* (upregulated in MB and in all but one PLEX sample in RNA-seq data and upregulated in MB in array data), carried EZH2 binding sites in brain and neural cells which were targeted by AT/RT-unique differentiation-related methylation (**Supplementary Table 4**). *DOK7* is shown to work in synaptic formation and *CAMK2B* in final steps of neural differentiation, making them potentially regulated by nBAF, as well (75,76). However, some of the genes linked to AT/RT-unique differentiation-related DMRs with EZH2 binding, such as *TNK1*, are found to be relevant in cancer development (77,78) and have not been associated with nBAF or late neural development. EZH2 binding sites were also detected in AT/RT-unique not-differentiation-related DMRs located in the promoter of AT/RT-specific DM-DE gene *TCEA3* (**Figure 3B-D, Supplementary Figure 11, Supplementary Table 4**).

Interestingly, the SWI/SNF subunits SMARCA4 and SMARCC1 were especially associated with DMRs that are AT/RT-unique and not differentiation-related (**Figure 4C**). SMARCA4, whose binding sites in other brain and neural cells were enriched in AT/RT hypermethylated DMRs (**Figure 2C**), was also enriched in sites that are AT/RT-specific and demethylated during neural differentiation (**Figure 4C**). These sites might represent hypermethylation of the neural SWI/SNF-EP300 target sites discussed earlier. Overall, TFs and regulators linked to SWI/SNF (8 out of 11) and H3K27ac (9 out of 11) were largely associated with AT/RT unique sites (**Figure 4C**).

Contrary to PRC2 and SWI/SNF subunits, a larger proportion (12 of 41, 29%) of neural TFs was enriched in PSC-like AT/RT hyper DMRs (**Figure 4C**), but PSC-like methylation was not linked to PRC2. Twelve TFs were enriched in both AT/RT unique and PSC-like regions. Notably, neural TFs BCHE and TFAP2A, the only neural TFs co-localizing with PRC2 subunits in AT/RT hypermethylated sites (**Figure 2D**), were also enriched only in AT/RT-unique AT/RT hyper-DMR categories (**Figure 4C**), linking them closely to PRC2-related DNA methylation.

In conclusion, our results suggest that AT/RT-specific methylation is linked to malfunctional SWI/SNF and PRC2 complexes. This aberrant DNA methylation has the potential to drive oncogenic epigenetic regulation, which is unique to AT/RT. In addition, we detected regions that harbor binding sites for several neural regulators and remained similar to PSCs that were originally methylated via a PRC2-independent mechanism.

### AT/RT-unique methylation in genes suppressed in AT/RT

To better understand the processes involved in AT/RT-related DNA hypermethylation, we more closely inspected the genomic neighborhoods of *CXXC5, TCEA3, NEUROG1, NEUROD2, IGSF9B* and *TNK1*. Typically, one DMR was detected in the gene loci with the used measurement techniques with the exception of *CXXC5* locus, which is located in a large-scale methylation area (in Chr5q31.2, **Figure 1G**) hypermethylated in ATRT when compared to MB and PLEX (**Figure 4D, Supplementary Figures 11 and 12, Supplementary Table 3**). It also harbors binding sites of several neural TFs and includes both PSC-like and AT/RT-unique DMRs. We originally discovered *CXXC5* to be downregulated in an AT/RT-specific manner in DM-DE gene analysis (**Figure 3B**) together with *TCEA3*, which carries AT/RT-unique, not-differentiation-related DMRs and neural EZH2 binding sites at the gene promoter. The binding sites of the SWI/SNF subunit SMARCA4 measured from rhabdoid tumor cells were detected in *CXXC5* and *IGSF9B*. In addition, EP300 binding sites measured from neural cells were also found around *CXXC5*, representing a possible neural SWI/SNF-EP300 target site described earlier (**Figure 4D**). EP300 binding sites in PSCs and brain tumors were also present in *IGSF9B* DMRs. *IGSF9B* and *TNK1* represent genes which were linked to AT/RT-unique methylation and include EZH2 binding sites. *TNK1* harbored binding sites of EZH2 and EP300 in neural cells, possibly representing an area that would normally be subjected to EP300 activity but is now methylated in AT/RT due to EZH2 (**Figure 4D**). TNK1 is expressed especially in progenitor cells and its loss leads to spontaneous tumor formation (77,78), suggesting a role as a tumor suppressor in less-differentiated cells.

Interestingly, both *NEUROG1* and *NEUROD2* were hypermethylated and downregulated in AT/RT when compared to MB in our DM-DE gene analysis and harbor DMRs, which are hypermethylated in AT/RT compared to PSCs and bound by BRD9 in rhabdoid tumors (**Figure 4D**). *NEUROG1* also includes EZH2 binding sites measured from several different cell types. However, the DNA-methylation difference was not significant in AT/RT-PLEX comparison most likely due to increased DNA methylation in PLEX AD subgroup (**Figure 3C, Supplementary Figure 10**), which includes more differentiated tumors than other tumor types and PLEX subgroups in our analysis. Both *NEUROG1* and *NEUROD2* were also lowly expressed in all PLEX subgroups (**Figure 3C, Supplementary Figure 10**) and are likely to be silenced in this differentiation lineage. *NEUROD2* DMRs showed AT/RT-unique DNA methylation but no reported EZH2 binding sites (**Figure 4D**). In MB, these two TFs behave in a complementary manner, as NEUROG1 is induced in all the other MB subgroups except in SHH subtypes, whereas NEUROD2 induction is more clearly detected in SHH and G4 subtypes (**Figure 3C, Supplementary Figure 10**). Overall, our results suggest that AT/RT-related DNA methylation is involved in suppressing these genes, which can contribute to impaired further neural development typical for AT/RT.

## Discussion

Our study aimed to identify DNA methylation patterns, transcriptional regulators associated with methylation differences, and aberrantly expressed genes specific to AT/RTs by using both array and sequencing data. AT/RTs are interesting in this respect, as the epigenetic oncogenic regulation presumably plays a key role in their oncogenesis. Other tumor types provided a good reference for AT/RT and allowed us to define AT/RT-specific features which also showed similar methylation patterns across different AT/RT subgroups. Our results consistently suggest that AT/RTs remain in a less differentiated epigenetic state compared to the other tumors, whereas the differentiation state is most advanced in MB with focal methylation of pluripotency-related TF target sites and demethylation-related activation of neural genes (79– 82). Although the characteristics of the materials, e.g. the genomic coverage, sample amounts, and suitable algorithms, differed, the key findings align.

We showed that the binding sites of several TFs that are the drivers of neural differentiation are hypermethylated in AT/RT, suggesting that TF activity in these sites is disrupted. NEUROG2 and NEUROD1 are especially notable examples because they are pioneer factors able to induce opening of the chromatin and to regulate a wide range of factors that affect neural differentiation (53,83–85). However, both belong to the basic helix–loop–helix (bHLH) gene family, members of which selectively bind to unmethylated DNA and are thus sensitive to DNA methylation (40). Furthermore, *e*.*g*. NEUROD1 target genes (83,86) OTX2 and EBF3, both with the highest expression in MB, have their binding sites also hypermethylated in AT/RTs, indicating the silencing of the whole differentiation cascade of genes. In our downstream analysis, *NEUROG1* and *NEUROD2* were also defined as DM-DE genes with higher methylation in AT/RT than PSCs, suggesting that they could serve as differentiation bottlenecks which are suppressed by DNA methylation in AT/RT. To summarize, NEUROG-NEUROD family TFs appear to be targets of DNA methylation-driven regulation at different levels, via decreased gene expression or altered DNA binding ability, both of which contribute to suppressed neural differentiation. Several other neural genes, such as *NBEA, BARHL1*, and AMER2 (87–89), were more methylated and less expressed in AT/RT than in MB. Our results are supported by prior notions that neuronal differentiation factors have lowered functionality in AT/RTs *e*.*g*. due to increased H3K27me3 near the genes or decreased gene expression in general (11,14,15,53,90–93). Our study sheds light to the relevant role of DNA methylation in their suppression.

Our results suggest that EZH2, the functional subunit of PRC2, contributes to AT/RT-specific hypermethylation. EZH2 inhibition suppresses an embryonic stem cell-like expression pattern in SMARCB1 deficient tumors (70), and EZH2 inactivation or deletion prevents AT/RT development and growth (15,70,94,95), making it a relevant regulator for these tumors. The main function of EZH2 is histone methylation leading to H3K27me3, which is also the main target of EZH2 inhibition, but decreased H3K27me3 levels suggest suboptimal or abnormal EZH2 activity in AT/RT (11). Instead, quiescent chromatin state, which lacks the measured chromatin marks, is increased in AT/RT and is likely to be targeted by DNA methylation (11). PRC2 marks many genes that are methylated during cell differentiation from PSCs to neural precursor cells (96). EZH2 has been reported to interact with DNA methyltransferases (96,97) and even to directly control DNA methylation (97), which could at least partly explain AT/RT-unique hypermethylation we observed. In rhabdoid tumor cells, DNA methylation is decreased and PRC2 members concomitantly released from the p16 locus upon SMARCB1 re-expression (98), also suggesting an interplay between PRC2 and DNA methylation.

When we further analyzed the EZH2 binding sites that harbor AT/RT-unique hypermethylation, we noticed that the genes linked to these sites often had functions related to late neural differentiation and nBAF, and part of them have been previously associated with cancer. Most of them were lowly expressed also in MB and/or PLEX, and are thus likely to be silenced by H3K27me3 or other factors. Although they are not linked to gene expression, they represent PRC2-associated DNA-methylation-mediated epigenetic regulation in AT/RT. However, we also detected EZH2 binding sites in AT/RT-unique DMRs in DE-genes like TCEA3 and TNK1, which are relevant for oncogenesis and early development, as well as AT/RT-unique DMRs lacking EZH2 binding sites in relevant genes, such as NEUROD2 (77,99). This might be due to incomplete EZH2 binding data (not all cell types are covered in GTRD) or because other factors are responsible for the differential DNA methylation.

Only a subpopulation of SMARCB1 deficient tumors responds to EZH2 inhibition in clinical trials (100,101), suggesting that the functional role of EZH2 is patient-specific and partly decreased after oncogenesis. Further studies have revealed marker genes and compensatory responses responsible for treatment resistance (102,103). The balance between EZH2-induced H3K27me3 and DNA methylation is still unclear, and constant EZH2 activity is most likely not needed for sustaining the DNA hypermethylation, which appears to have a significant role in AT/RT. On the other hand, most of the EZH2-associated genes were linked to later stages of neural differentiation and not differentially expressed between tumor types, questioning the role of EZH2-mediated DNA methylation in AT/RT malignancy.

PSC-like DNA methylation and gene expression patterns have been shown to exist in human AT/RT samples (70). We showed that areas similarly methylated in both AT/RTs and PSCs are less methylated in MB and PLEX and differentiation-related, giving them the function and strengthening their relevance. This finding suggests that these PSC-like sites remain methylated, and thus inaccessible, in AT/RT, which maintains AT/RTs in a low differentiation state and promotes malignancy. We also showed that many of these regions involve large-scale methylation differences, implying larger changes in the chromatin structure. What is the relationship between these PSC-like sites and AT/RT-specific sites that do not co-localize with the binding sites of most neural TFs? SWI/SNF complex has been shown to guide demethylation during cell differentiation (104–106) Thus, the PSC-like methylation can reflect SWI/SNF complex’s PRC2-independent disability to target demethylation in the absence of SMARCB1. AT/RT-unique DNA methylation can also halt AT/RT cells at a specific point in the developmental trajectory by suppressing the expression of key neural regulators, such as NEUROG1 and NEUROD2, and in this way prohibit the downstream demethylation steps necessary for further neural development. It is likely that at least part of PSC-like methylation is due to these downstream effects, as SMARCA4 binding is enriched in AT/RT unique sites and has not been detected in many PSC-like regions.

In summary, our analysis revealed DNA methylation patterns specific to AT/RT and associated them with gene regulation, cell differentiation, and malignancy. AT/RTs carry very few genetic alterations, all of them targeting the SWI/SNF epigenetic regulatory complex; therefore, aberrant epigenetic regulation is relevant for these tumors. Since other studied epigenetic marks are mainly depleted in AT/RTs (11), the elevated DNA methylation we observed is likely to play a role in AT/RT malignancy. Our results suggest that targeted DNA demethylation is one of the key processes during neural cell differentiation, which exposes the enhancers and promoters of neural genes for controlled gene activation. In AT/RT, this process has not taken place, and DNA methylation is also targeted to novel sites. AT/RT-unique DNA methylation also appears to silence key neural regulators and other neural genes. It is known that SMARCB1 loss needs to happen during certain developmental stages for rhabdoid tumor development, whereas mainly lymphomas and benign tumors are formed at later phases (107–109). Based on our results, the temporal relationship to the DNA demethylation might be one of the key determinants of the cell fate after SMARCB1 inactivation. Further research will show whether treatment options affecting DNA methylation and differentiation blockage could generate positive outcomes for patients suffering from this devastating disease.

## Materials and Methods

### Sequencing cohort

Tumor samples analyzed with next-generation sequencing were collected for this study from 10 patients operated at Tampere University Hospital. The study was performed in line with the principles of the Declaration of Helsinki. Approval for this study was granted by The Regional Ethics Committee of Tampere University Hospital (decisions: R13050, date 9.4.2013, R14024, date 28.3.2014, and R07042, date 20.9.2017) and Valvira (decision V/78697/2017). An experienced neuropathologist evaluated the tumor samples and determined the histopathological type and grade according to the criteria presented by the WHO 2016 (110).

### Reduced representation bisulfite sequencing

DNA was isolated from the frozen samples using a QIAamp DNA Mini kit (QIAGEN) with RNAse treatment and from the FFPE sample with a turXTRAC FFPE DNA kit (Covaris). Library construction and sequencing were performed at the Beijing Genomics Institute (BGI), Hong Kong. Samples were sequenced using Illumina HiSeq2000 technology. Paired-end sequencing of 50 bp was used. The data goal was 5Gb.

### RNA sequencing

RNA was isolated using a mirVana isolation kit (Invitrogen). Library construction and sequencing were performed at Novogene, Hong Kong. Samples were sequenced using Illumina HiSeq2000 technology. Paired-end sequencing of 150 bp was used. The data goal was 20 million raw reads per sample.

### Preprocessing and analysis of DNA methylation datasets

#### i450k data

The full CNS tumor reference cohort GSE90496 established by Capper et al. was downloaded from Gene Expression Omnibus (GEO) (111,112), filtered, batch-effect corrected, and normalized as described in their study (26), resulting in an initial set of 428799 probes and 2801 samples. Capper method includes the following probe filters: 1) removal of probes targeting the sex chromosomes, 2) removal of probes containing a single-nucleotide polymorphism (dbSNP132 Common) within five base pairs of and including the targeted CpG site, 3) probes not mapping uniquely to the human reference genome (hg19) allowing for one mismatch, and 4) probes not included on the Illumina EPIC array. Supplementary clinical annotations were also mapped onto the samples. AT/RT, MB, PLEX, and CONTR samples, excluding the control subtype “INFLAM” and all samples without location information, were then extracted (N = 584).

To detect DMRs between tumor types, AT/RT-MB, AT/RT-PLEX, and PLEX-MB comparisons were conducted, and each cancer was also tested against the CONTR set. Additional probe filtering (removing probes not mapping to hg38 genome and the cross-hybridizing probes) followed by DMR calling for the remaining 396700 probes was performed with DMRcate v1.18.0, switching the hg19 coordinates into hg38 with rtracklayer v1.42.2 (113–115). The beta cutoff-threshold was set to 0.05, and FDR < 0.05 but otherwise the default settings were used. Location information was included in the linear model to adjust for biological confounders. To reduce normal cell effects, the DMRs of each main comparison (*e*.*g*., AT/RT-MB) was overlapped with the control DMRs (*e*.*g*., AT/RT-CONTR, MB-CONTR) using subsetByOverlaps-function. The resulting regions were reduced by taking only the intersecting regions between the main comparison and at least one of the control comparisons (116). Based on the visual inspection of the regions using heatmaps, the final differential methylation criteria used to define the DMRs most relevant to further analyses was increased from 0.05 to 0.20.

DMRs were annotated to annotatr v1.8.0 pre-built hg38 genic regions, CpG regions, and FANTOM5 enhancers (117,118). In addition, the pre-built promoter annotations (1 kb upstream from the transcription start site [TSS]) were used to generate broader promoters (2 kb upstream and 500 bp downstream from the TSS), and genomic neighborhoods spanning 200 kb in both directions from the TSS.

#### RRBS data

Quality control, adapter trimming, and MspI restriction site trimming were performed using the Babraham Bioinformatics Group preprocessing tools FastQC v0.11.7 and Trim Galore v0.5.0. Reads were queried against bisulfite-converted reference genomes of potential contaminants with FastQ Screen v0.13.0 to check the origin of the libraries (119). Mapping against hg38 (UCSC) reference and mapping quality control without duplicate removal was performed using Bismark v0.19.1 (120). The quality information of all these steps was summarized over the dataset using MultiQC (121).

Methylation percentage calling and differential methylation analysis were performed using methylKit v1.8.1 (115,122). CpGs with coverage of less than 10 reads or hitting into scaffolds, sex chromosomes, or mitochondrial DNA were removed. The CpGs were tiled into 1000 bp non-overlapping regions, i.e., tiles, which were used for DMR calling with overdispersion correction. Regions with methylation percentage difference greater than 25% and q value (FDR) smaller than 0.05, were extracted and then annotated similarly as i450k data.

#### Dimensionality Reduction

The beta values for all AT/RT, MB, and PLEXUS samples were aggregated on RRBS tiles based on lift-over hg38 probe coordinates. This step was performed by taking the complete overlaps and then calculating the mean beta value for probes hitting each tile (116). The scales in the dataset were unified, and the 10 000 most varying regions were used as inputs for various dimensionality reduction algorithms.

#### K-means clustering

The i450K DMRs (q<0.05, meth. fold change >= 0.20, three comparisons) were combined into a single pool of regions (n=3780). These were used to generate a methylation matrix such that the mean of the probes hitting each region was used to represent the methylation level for the region in each sample. K-means clustering was calculated with the cluster amounts from 1 to 25, each time using 25 iterations and 50 starting points. The optimal number of clusters was estimated with Akaike information criterion (AIC), resulting in k=10. DMR behavior in clusters was visually inspected using heatmaps. Moreover, the regions belonging to each cluster were aggregated using the median to summarize the average methylation patterns in the sample types.

### TF binding analyses

#### Preliminary Statistical Testing

TF binding site (TFBS) enrichment analysis was performed using one-sided Fisher’s exact test (FET). TFBS data originating from ChIP-seq measurements were obtained from GTRD v.19.04 metaclusters (39). Before testing, DMRs with beta fold change >= 0.25 were filtered by overlap operations between the comparisons to obtain the regions specific to each cancer with a concordant direction of methylation change. Hypermethylated and hypomethylated regions were tested separately. For the i450k data, the background set was generated from previously filtered 396700 probes, and for the RRBS data, the methylKit tiles were used as the background.

When performing the tests with i450k data, the hg38 coordinates of each 50 bp probe overlapping with DMRs were first obtained. These were extended to 500 bp in both directions, and the methylation fold-change of the original DMR was used to represent the methylation difference in each extended probe. Regions were considered cancer-specific if they overlapped with at least 600 bp between comparisons, and with RRBS, because of the non-overlapping nature of the DMRs, the exact overlaps could be used.

FET p-values were adjusted by Benjamini-Hochberg (BH) correction (123). TFs with adjusted p-values smaller than 0.05, were considered significant. Sets of validated TFs with significant enrichment in the analogous DMR-sets, that is, in hypermethylated or hypomethylated cancer regions of the i450k and RRBS experiments, were constructed. Validated TFs had been studied in the literature, and because common themes existed, they were manually annotated into 11 selected categories.

TFBS enrichment analysis to k-means clustering results was carried out using a similar method to the above (One-sided Fisher’s exact test, BH correction). Cancer specific DMRs from 450k data were assigned to the clusters and TFBS enrichment was tested for all GTRD TFs with all clusters, cancer types and methylation changes (hyper/hypo) by examining all autosomal 450k probes and associating them with TF binding and DMRs to create the contingency table.

#### Customized TFBS enrichment analysis with cell-type splitting

GTRD annotations for cell types linked to a given TFBS were further examined and manually divided into 11 groups (**Supplementary Table 2**). The GTRD was then split based on the terms of the groups, and these groups were used as input material for the FET to replace the full GTRD.

#### Studying average methylation levels at binding sites

Methylation of the relevant i450k probes near the binding sites of the validated TFs (maximum gap 500 bp) was visualized as violin plots. The probes were selected by pooling the cancer regions and then finding complete overlaps with the hg38 probe coordinates. The beta values over each tumor subclass were aggregated probewise using the mean.

#### TFBS co-localization

The co-localization of validated TFs was studied with TF binding sites hitting target areas (hypermethylated AT/RT regions, hypermethylated MB regions, hypomethylated MB regions, or hypomethylated PLEX regions). TFs were tested pairwise against each other to construct four-field matrices for each case. Regions with 1) no binding site from either TF, 2) binding sites from both TFs, and 3-4) binding sites from one TF but not the other were counted and the significance was tested using one-sided FET and stored as a symmetric TF matrix of P values. These P value matrices were corrected using the BH method. Adjusted P values were subsetted by the TF names of the validation group (AT/RThyper-TFs, MBhyper-TFs, MB-hypoTFs, PLEXhypoTFs) and visualized using the ComplexHeatmap package (124).

### Differential expression analysis for expression datasets and integration with methylation data

#### Array datasets

Normalized expression arrays of three public datasets were downloaded from GEO: IIlumina HumanHT-12 V3.0 (125) (accession GSE42658) and two Affymetrix Human Genome U133 Plus 2.0 sets (126) (accession GSE35493 and (127), accession GSE73038). The Illumina set contains measurements of all three cancers, and Affymetrix sets provide additional support for AT/RT-MB comparison. Before DE analysis with limma, the expression matrices were log2-transformed if that had not been performed by the normalization method (128). Probes were annotated into symbolic gene names, and the mean of all the probes mapped to the same gene was chosen to represent the expression values. Genes with BH adjusted p values smaller than 0.05 and log2-transformed fold changes (absolute change) of at least 1 were considered significant.

#### RNA-seq data

After sequencing data QC with FastQC v0.11.7 (https://www.bioinformatics.babraham.ac.uk), reads were mapped and quantified against hg38 by kallisto v0.44.0 (129). The kallisto transcript abundances were transformed to gene level counts and scaled to library size using tximport v.1.10.1. DE analysis was then performed using DESeq2 v1.22.2, with default settings (130). Genes with BH adjusted p-values smaller than 0.05, and log2-transformed fold changes (absolute change) of at least 1 were considered significant.

#### Integration

The DE results were merged with the annotated DMRs by symbolic gene names to find cases where the direction of the methylation change in a regulatory region (i.e., promoter, enhancer, genomic neighborhood) was opposite to the gene expression change. Similar analysis was also done for the genes with concordant up/downregulation of both DNA methylation and gene expression. GEO expression array results were integrated with i450k DMRs, and RNA-seq based results were integrated with RRBS DMRs from matched cases to provide data for validation. Validation was performed by extracting the genes that showed similar behavior in both approaches. Since annotatr uses transcript-level annotations, the validated hits, *e*.*g*., for promoters, represent DE genes that show regulatory methylation in at least one of its transcript specific promoters in both datasets. Additional filtering and validation steps were performed for the enhancers and genomic neighborhoods, described later. The results of the experiments were summarized using Venn diagrams and an oncoprint-like visualization using the ComplexHeatmap package (124).

A recent regulatory genomics workflow was adapted to integrate DMRs and DE genes with precalculated promoter-enhancer annotations provided by the FANTOM5 database (118,131). DE-DMR pairs in genomic neighborhoods were queried between i450k and RRBS experiments with a maximum gap of 5 kb, and the regions inside this “validation distance” with similar methylation direction were extracted. Moreover, to ensure that a given DMR can spatially interact with the TSS of a given gene, the results were studied with TAD information from Hi-C experiments (132,133) (http://3dgenome.fsm.northwestern.edu/). The region between the DMR and TSS of each record overlapped with TADs originating from five cell types: SKNDZ, Cortex DLPFC, Hippocampus, H1-ESC, and H1-NPC (132). Genes without any TAD boundaries in any of their transcript TSS-DMR pairs were extracted.

### Linking DMRs to TADs for large-scale DNA methylation analysis

DMRs were overlapped with *Cortex DLPFC* and H1-NPC TADs by using Bedtools GroupBy v2.25.0 and in-house scripts to calculate hyper- and hypomethylated records in each TAD (132,134). The input material was either a broadened (+-500bp) probe extracted from the i450k DMRs or the methylKit DMRs of the RRBS analysis. The resulting TADs for each comparison were filtered such that at least five similarly directed probes/tiles, each with the methylation difference of at least 25 % with both the i450k and RRBS approach, should hit the TAD. Simultaneously, less than 1/10 of the hits can undergo methylation changes in the opposite direction. Moreover, the distance between the first and last probe or tile in the TAD had to be over 50 kb. The filtered TADs from different cell lines were pooled and reduced to produce single TAD coordinates for each comparison and then visualized with karyoploteR (135).

### Comparing brain tumors to ESCs and primary fetal brain

Two PSC samples, nine primary FB samples (GSE116754) (72), and 25 PSC samples (GSE31848) (71), were downloaded from GEO and preprocessed using the Capper et al. data (26). Beta-value density plots and tSNE-clustering for the 10 000 most varying probes were used to check that samples were separated based on tumor and normal sample type. The DMRs were called with DMRcate by comparing AT/RT, MB, and PLEX samples against PSC and FB cells. In addition, PSC samples were compared with FBs.

From these seven comparisons, hypermethylated and hypomethylated DMRs were extracted (q < 0.05, abs. average beta fold change >= 0.25 and length between 100 and 5000 bp). The region lists were overlapped with the same cancer-specific regions extracted for TFBS analysis (min 1 bp overlap). As a result, the experiment was summarized in a “developmental DMR matrix” of 2301 non-overlapping regions showing which of the comparisons or cancers each region overlapped. The resulting region sets were categorized based on the differences detected in tumor-tumor, tumor-normal, and PSC-FB comparisons. TFBS enrichment analysis was performed for these categories by extracting and extending the probes hitting the regions and testing them against GTRD using one-sided FET, as described.

## Ethic statement

The use of clinical material was approved by the ethical committee of the Tampere University Hospital (approval numbers R13050, R07042 and R14024).

## Acknowledgements

We would like to acknowledge Mrs. Paula Kosonen, Mrs. Päivi Martikainen, Mrs. Marika Vähä-Jaakkola, Mrs. Marja Pirinen, Mrs. Sari Toivola, Mrs. Hanna Selin, Mrs Marita Nieminen, Mrs Leena Jalonen and Mrs Satu Ranta for sample handling and logistics, Mrs. Aliisa Tiihonen and Mrs Maria Pere for scientific input, Mrs. Suvi Lehtipuro for preprocessing of i450k data, and Mr. Tomi Häkkinen for technical support. Personnel at Tampere University Hospital and Fimlab laboratories Ltd. are acknowledged for their contribution to sample collection. We are grateful for them and the patients for permitting the analysis of patient material. The study was financially supported by the Academy of Finland (#312043 (MN), #310829 (MN), #333545 (KG)), Cancer Foundation Finland (MN, KG), Sigrid Jusélius Foundation (MN, KG), Emil Aaltonen Foundation (KG), Finnish Cancer Institute (MN), Competitive State Research Financing of the Expert Responsibility area of Tampere University Hospital (MN, KG), Väre Research foundation (KN), The Finnish Pediatric Research Foundation (KN) and Finnish Cultural Foundation (JU). We acknowledge the CSC - IT Centre for Science, Finland, for providing computational resources and the Biocenter Finland (BF) and Tampere Genomics Facility for the service.

