## Supplemental Figures and Legends for "Aberrant DNA methylation distorts developmental trajectories in atypical teratoid/rhabdoid tumors"

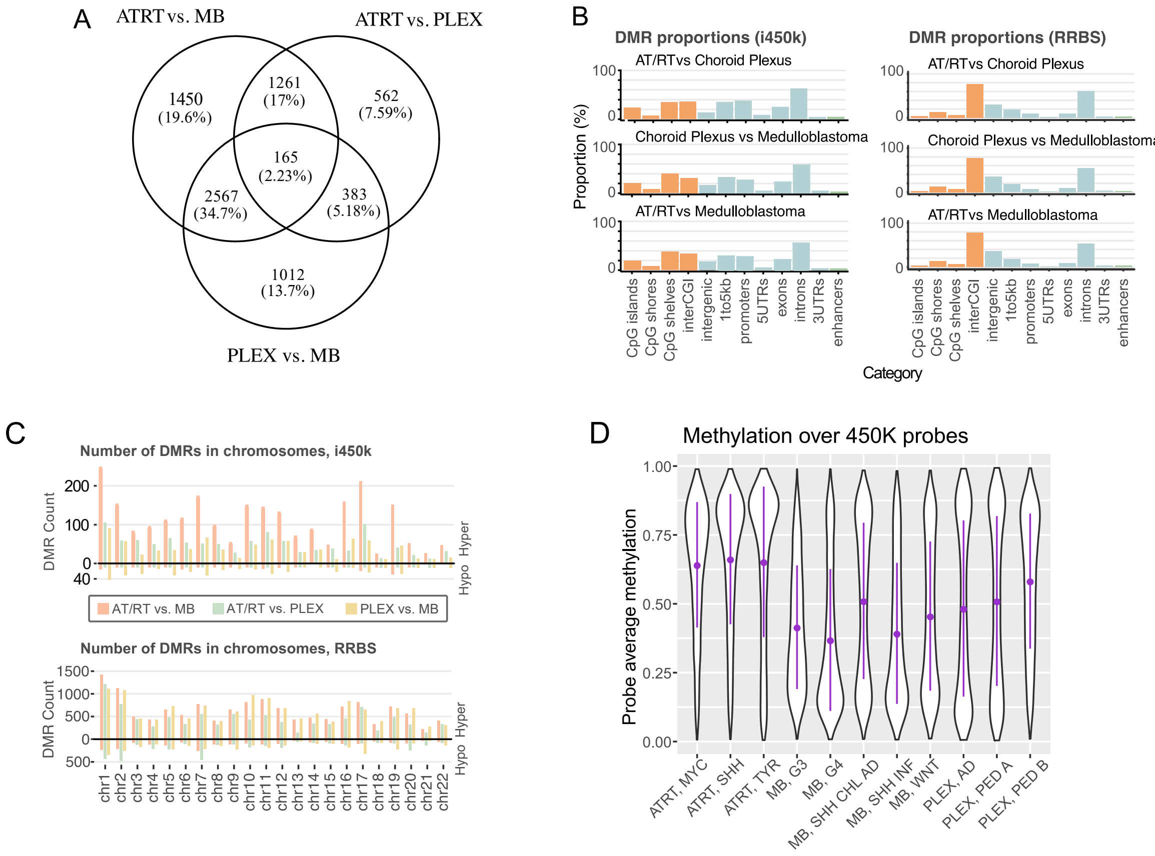


**Supplementary Figure 1**

**A.** Venn diagram of i450k DMRs obtained without filtering based on tumor location or normal tissue samples. **B.** DMRs in different comparisons show similar patterns, but a higher proportion of DMRs was covering interCGI and intergenic regions in RRBSdata than in i450k data. Normalized proportions of DMRs hitting to annotatr built-in CpG island annotations (in orange), genomic annotations and enhancers (in green) are visualized 94. The i450k and RRBS results were made comparable by annotating each bp from DMRs separately and normalizing the annotation counts with the total length of DMRs in each comparison. InterCGI means areas not hitting to CpG islands, shelves or shores, i.e. the "open sea". 1to5kb contains regions 1 to 5kb upstream of TTS. **C.** The distribution of DMRs into autosomal chromosomes. The majority of DMRs are hypermethylated in AT/RT and hypomethylated in MB when compared to other tumor types. **D.** Methylation distribution in i450K probes in AT/RT, MB, and PLEX subgroups . Violin plots visualize the distribution of DNA methylation in the 10 000 most variable sites in i450k microarray dataset.

**Supplementary Figure 2**

A

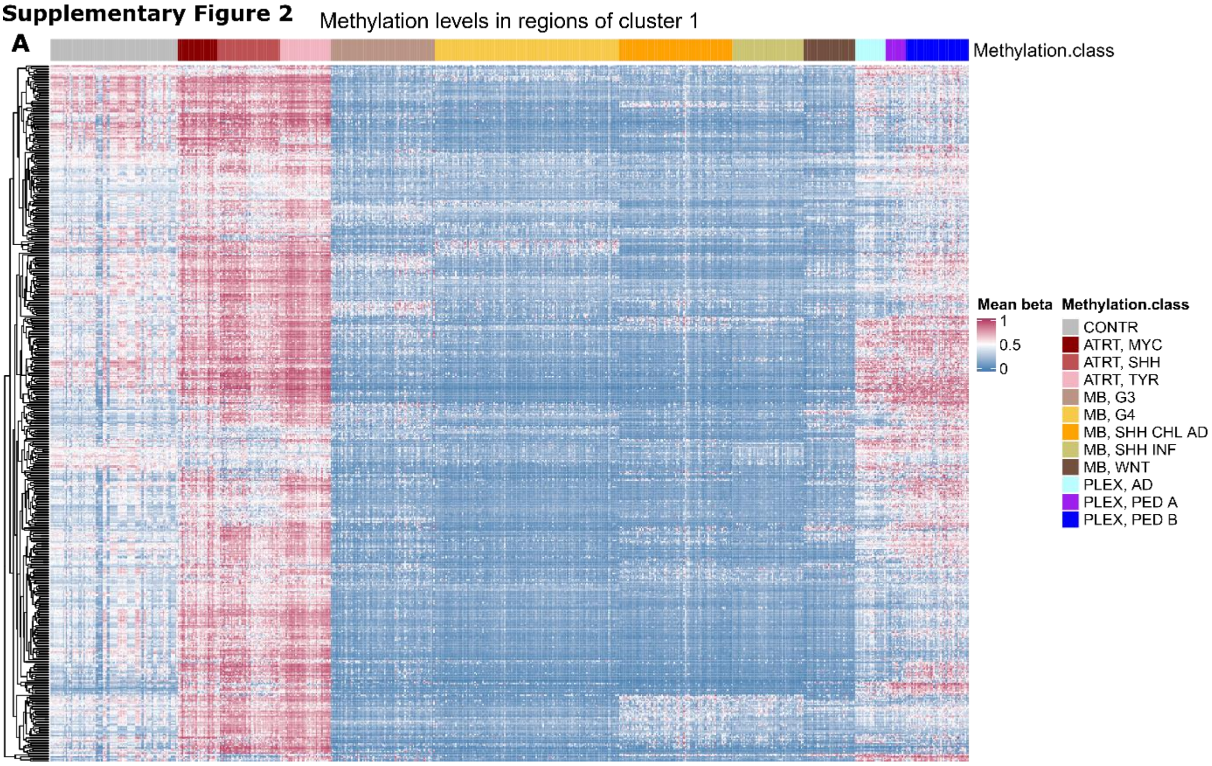


B

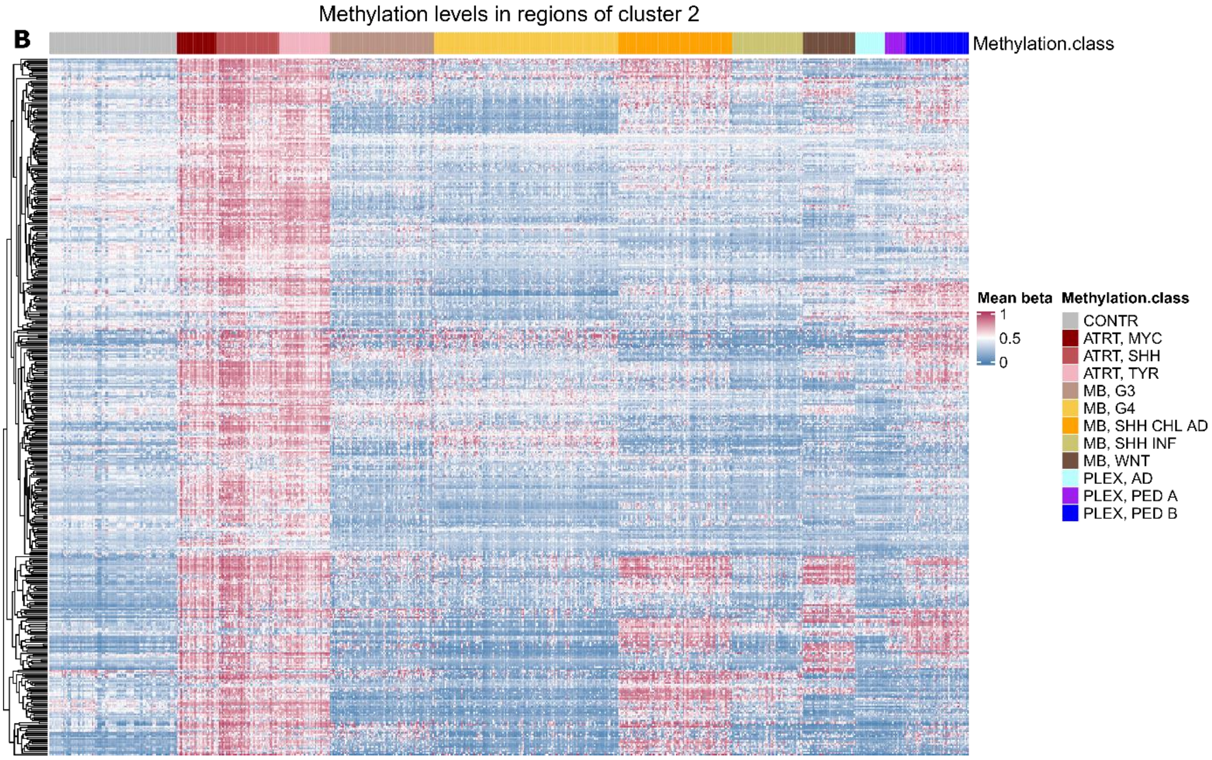

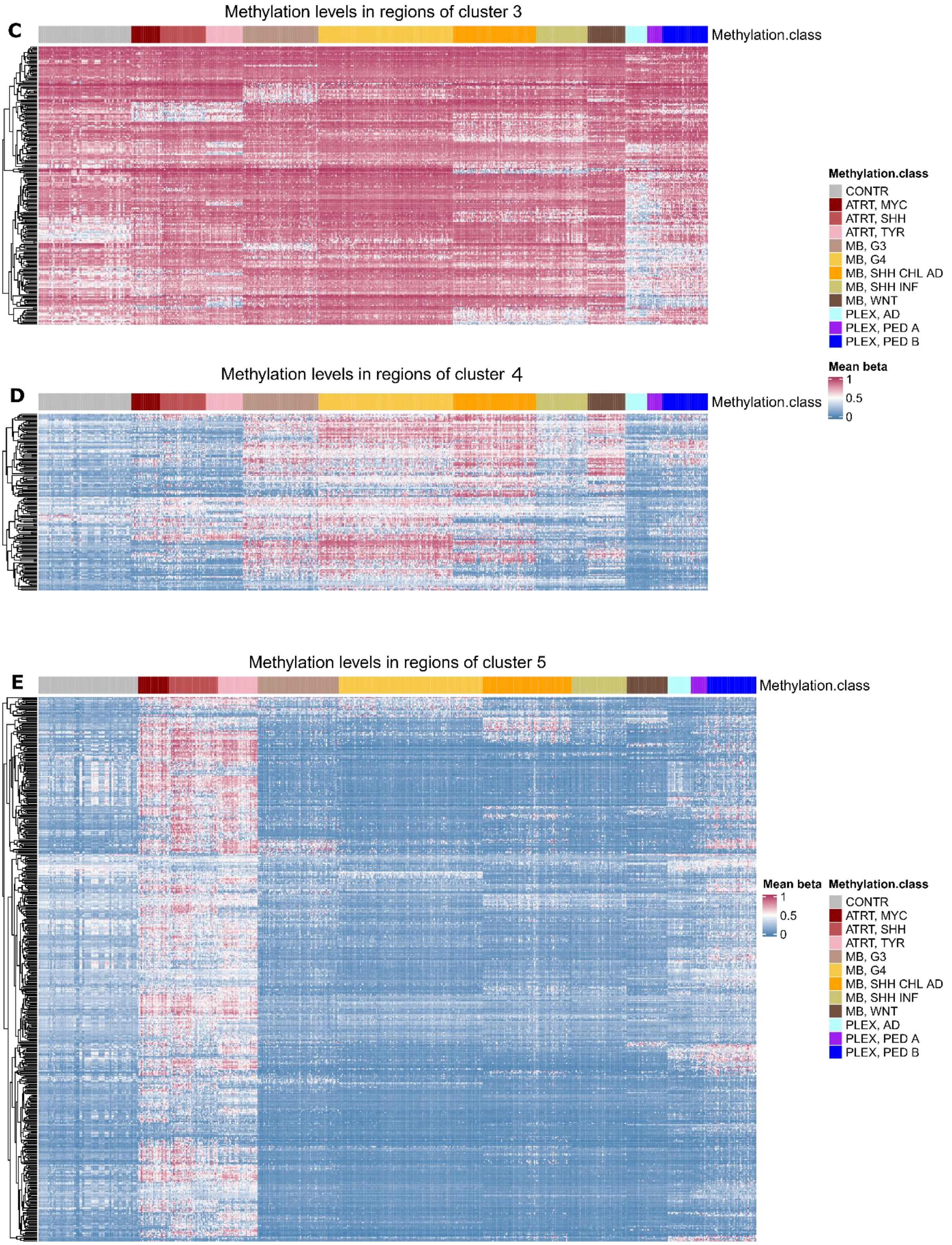
Methylation levels in regions of cluster 3
C
Methylation.class
Methylation.class
CONTR
ATRT, MYC
ATRT, SHH
ATRT, TYR
MB, G3
MB, G4
MB, SHH CHL AD
MB, SHH INF
MB, WNT
PLEX, AD
PLEX, PED A
PLEX, PED B
Mean beta
1
0.5
0
Methylation levels in regions of cluster 4
D
Methylation.class
Methylation levels in regions of cluster 5
E
Methylation.class
Mean beta
1
0.5
0
Methylation.class
CONTR
ATRT, MYC
ATRT, SHH
ATRT, TYR
MB, G3
MB, G4
MB, SHH CHL AD
MB, SHH INF
MB, WNT
PLEX, AD
PLEX, PED A
PLEX, PED B

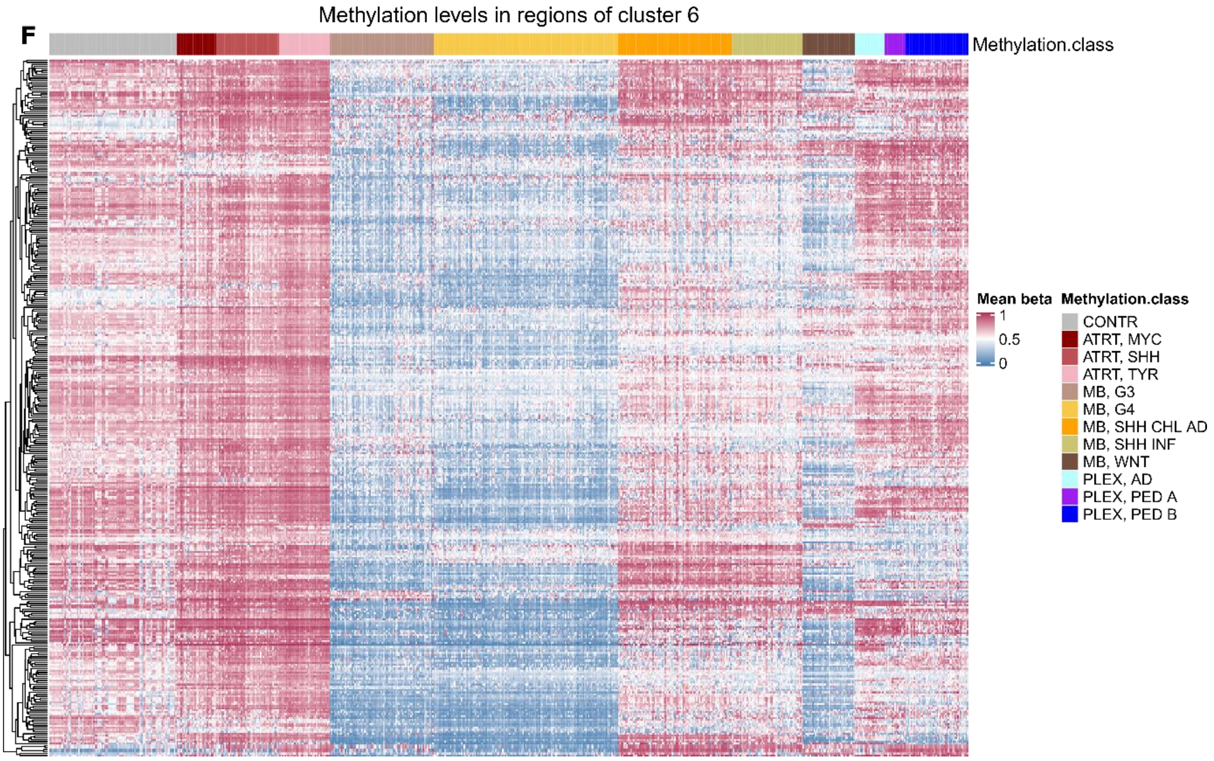
F
Methylation levels in regions of cluster 6
Methylation.class
Mean beta
1
0.5
0
Methylation.class
CONTR
ATRT, MYC
ATRT, SHH
ATRT, TYR
MB, G3
MB, G4
MB, SHH CHL AD
MB, SHH INF
MB, WNT
PLEX, AD
PLEX, PED A
PLEX, PED B

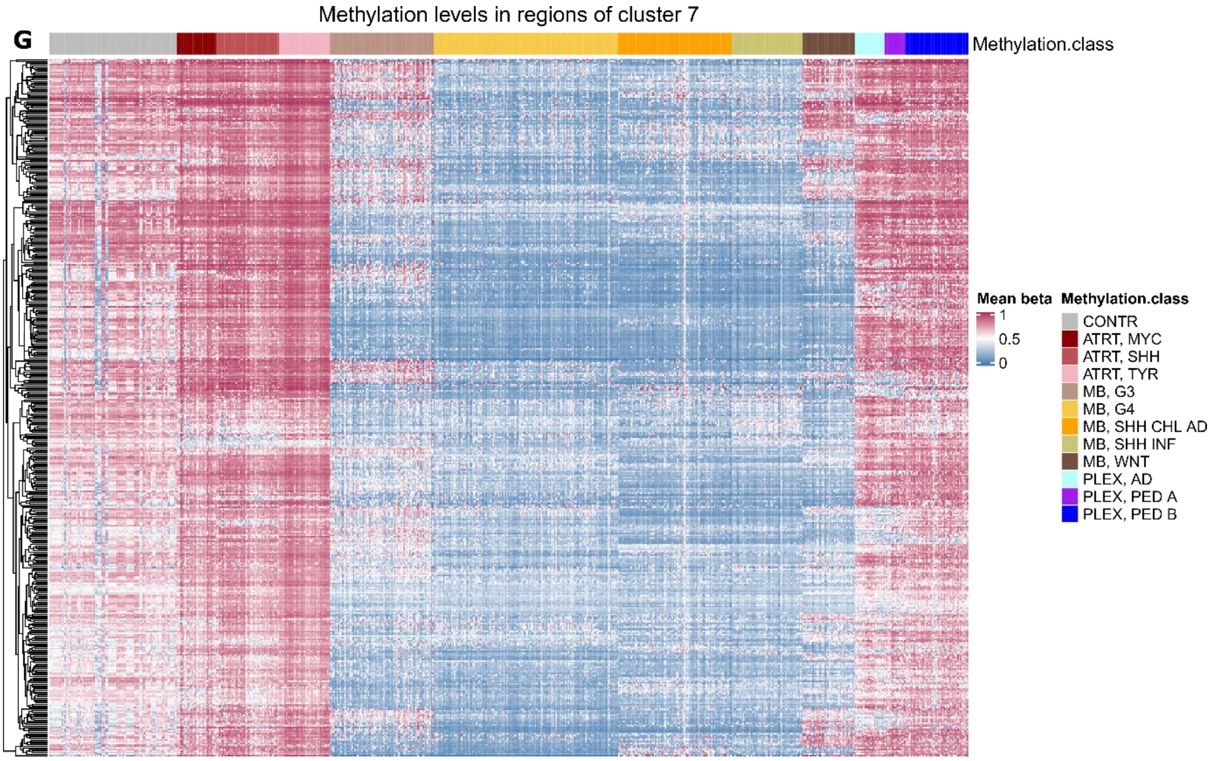
G
Methylation levels in regions of cluster 7
Methylation.class
Mean beta
1
0.5
0
Methylation.class
CONTR
ATRT, MYC
ATRT, SHH
ATRT, TYR
MB, G3
MB, G4
MB, SHH CHL AD
MB, SHH INF
MB, WNT
PLEX, AD
PLEX, PED A
PLEX, PED B

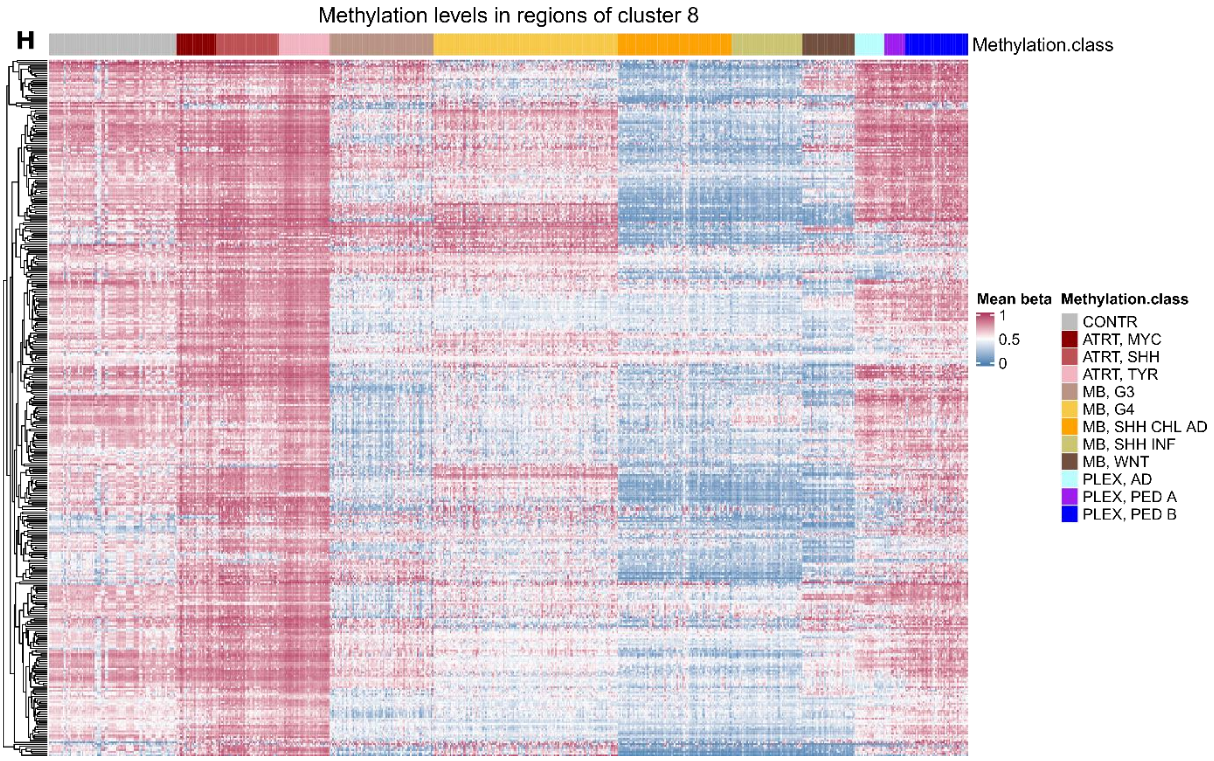
Methylation levels in regions of cluster 8
H
Methylation.class
Mean beta
1
0.5
0
Methylation.class
CONTR
ATRT, MYC
ATRT, SHH
ATRT, TYR
MB, G3
MB, G4
MB, SHH CHL AD
MB, SHH INF
MB, WNT
PLEX, AD
PLEX, PED A
PLEX, PED B

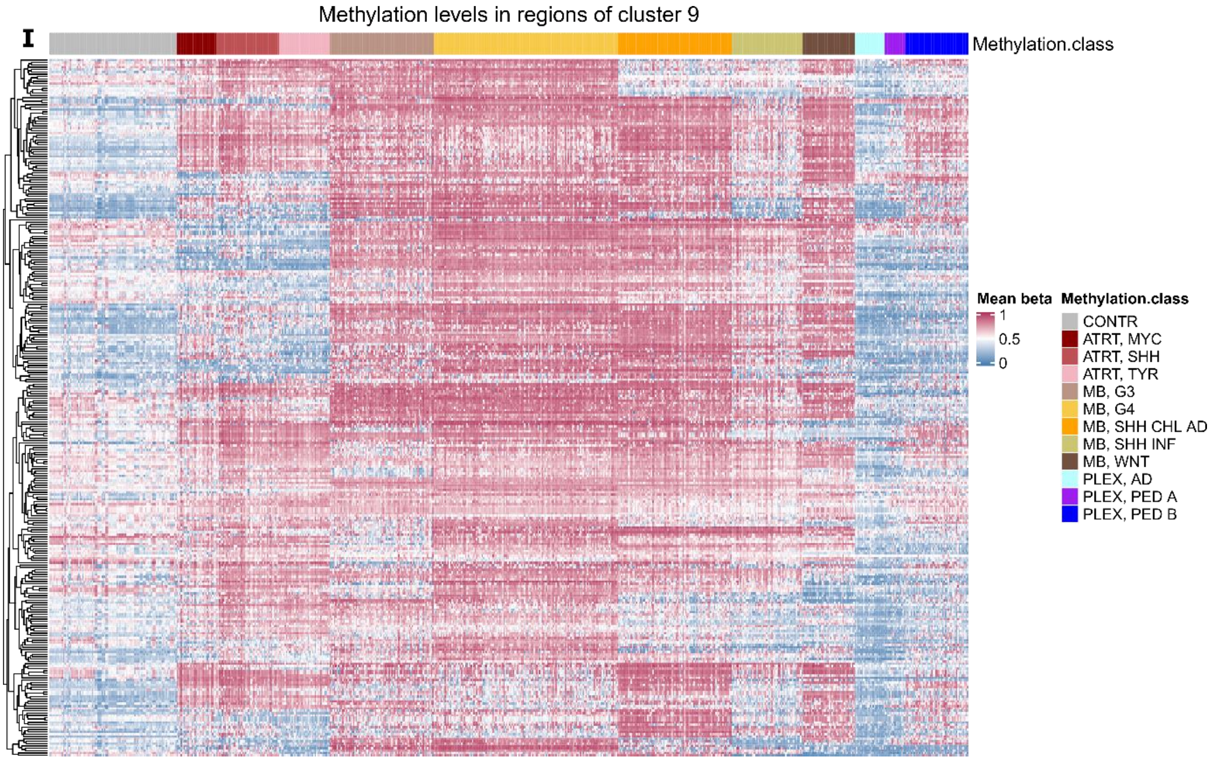
Methylation levels in regions of cluster 9
I
Methylation.class
Mean beta
1
0.5
0
Methylation.class
CONTR
ATRT, MYC
ATRT, SHH
ATRT, TYR
MB, G3
MB, G4
MB, SHH CHL AD
MB, SHH INF
MB, WNT
PLEX, AD
PLEX, PED A
PLEX, PED B

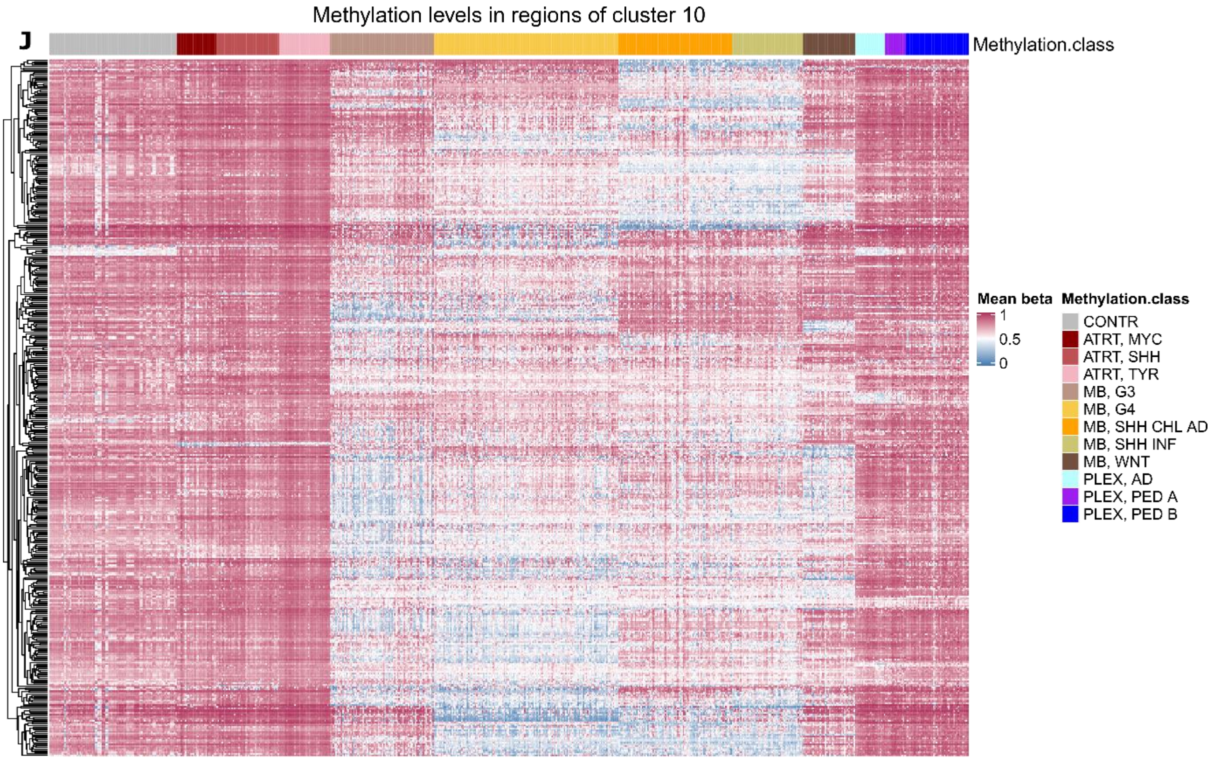


**Supplementary Figure 2**
A-J. Corresponding cluster heatmaps for k-mean clustering. Methylation class shows the tumor type and subtype.

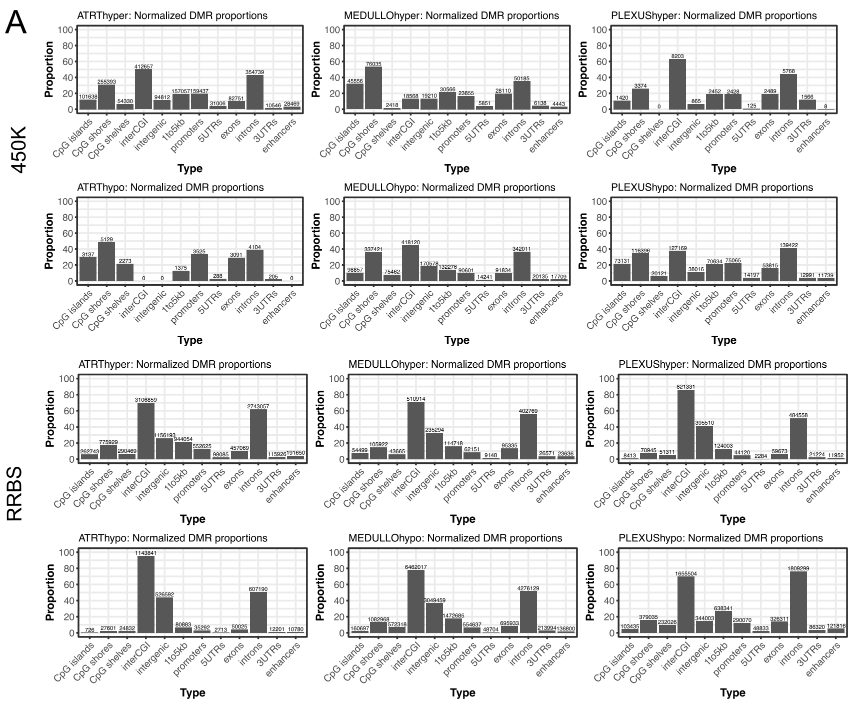


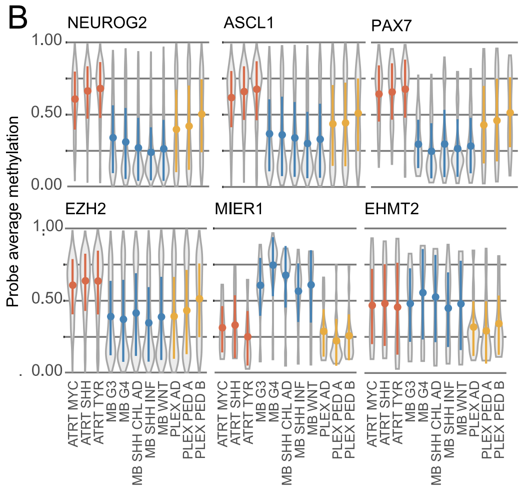


**Supplementary Figure 3**

**A.** Genomic annotations for cancer specific DMRs in all tumors for RRBS and i450k data. A higher proportion of MB hypermethylated DMRs (7.6% and 32% in RRBS and i450k data, respectively) w ere located in CpG islands, w hen compared to MB hypomethylated, AT/RT hypermethylated or PLEX hypomethylated DMRs (1.9-5.9% and 11-12% in RRBS and i450k, respectively). **B.** Binding sites of neural differentiation factors NEUROG2, ASCL1 and PAX7 are more methylated in AT/RT, irrespective of the tumor subtype. Out of TFs linked to histone lysine methylation, EZH2 (involved in histone H3 lysine 27 trimethylation) behaved similarly, but a distinct DNA methylation pattern w as observed for MIER1 and EHMT2 (involved in histone H3 lysine 9 methylation) binding sites. Violin plots visualizing the distribution of DNA methylation w ith selected TF binding sites in all i450k DMRs. The tumor subgroups are presented separately.

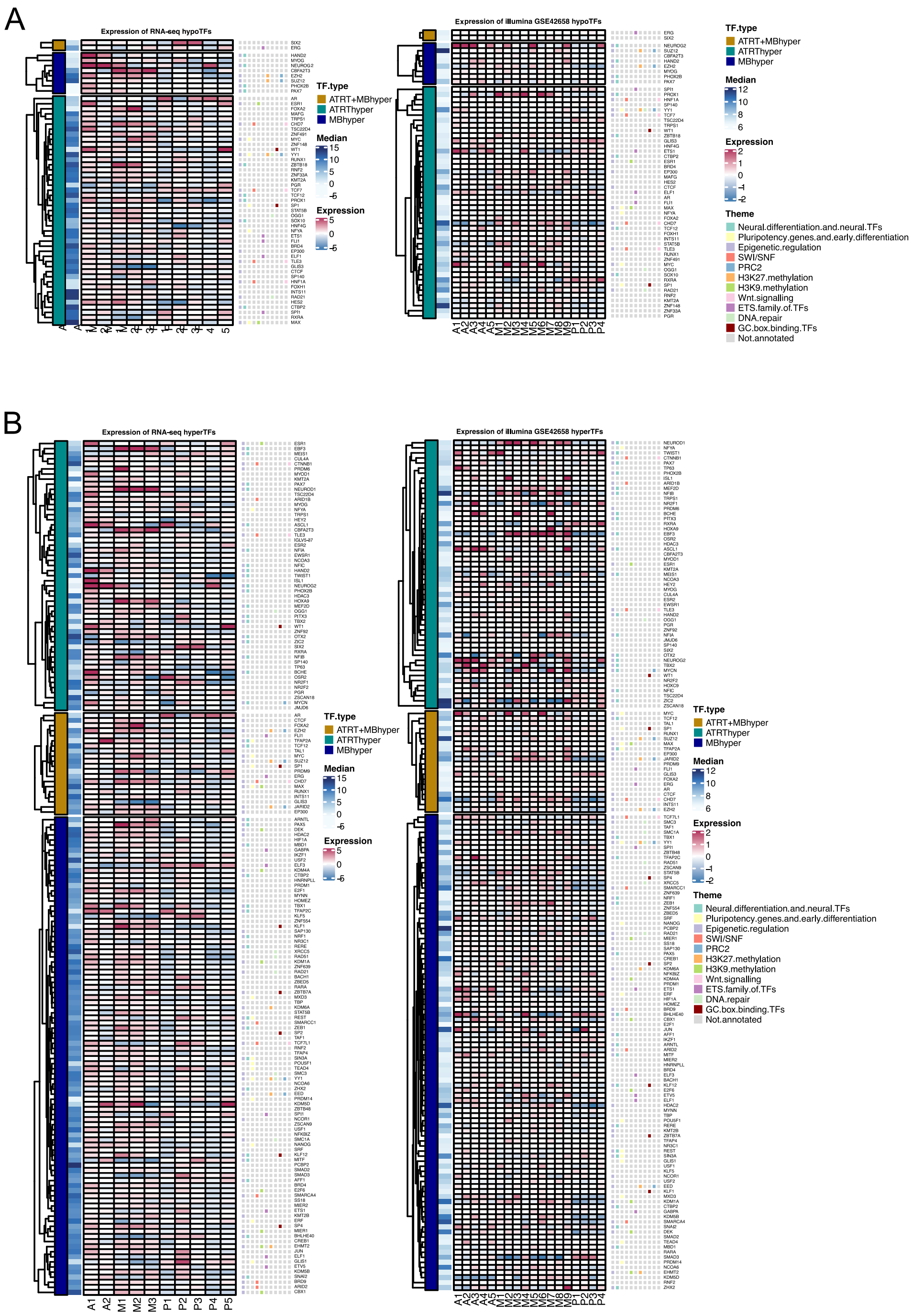


**Supplementary Figure 4.**

**A.** Heatmaps visualizing the expression of TFs associated with tumor type spesific hypomethylated DMRs in RNA-seq and microarray data. **B.** Same as in A but for hypermethylated DMRs.

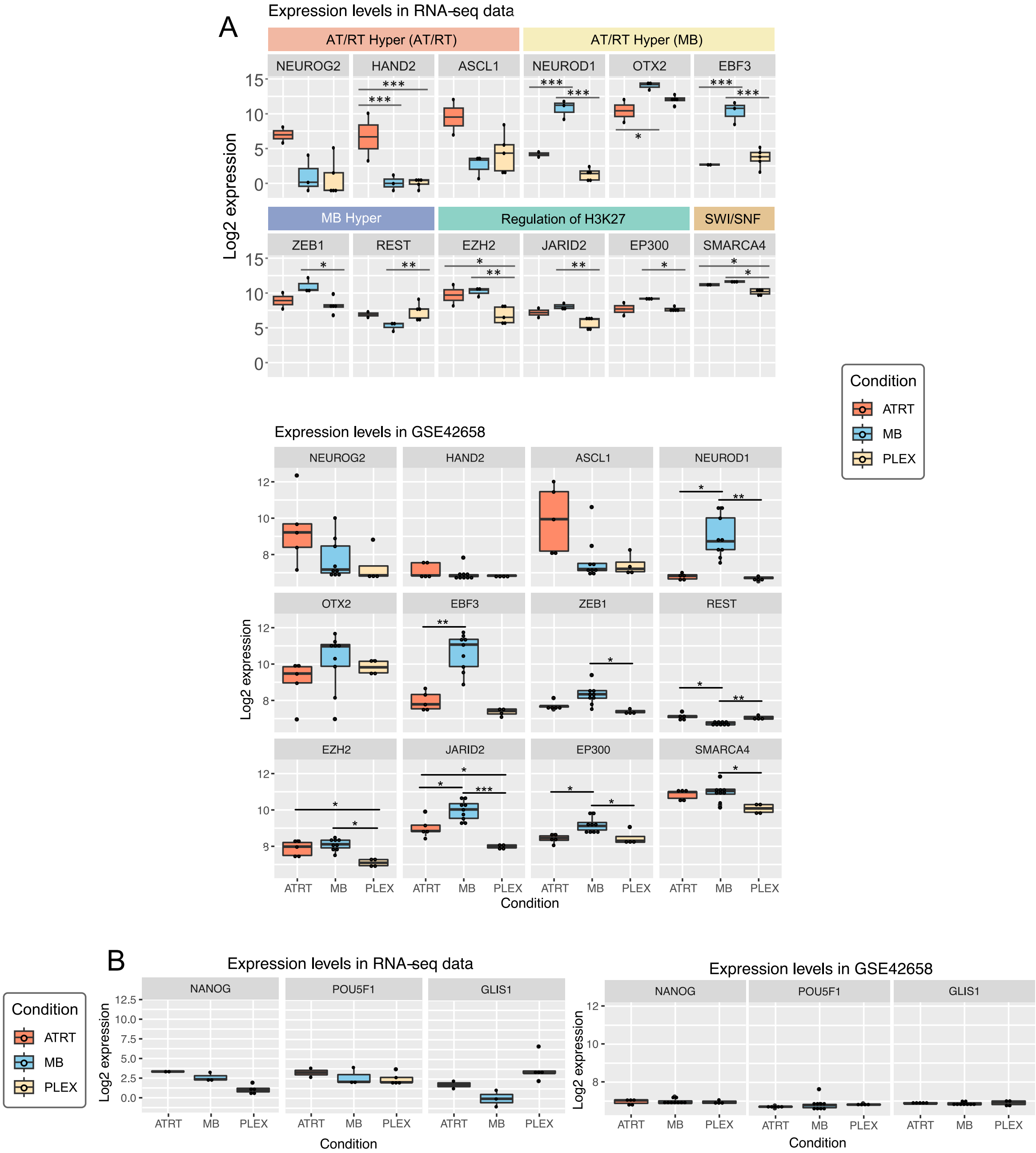


**Supplementary Figure 5**

**A.** Neural dfferentiation related TFs enriched in DMRs hypermethylated in AT/RT (AT/RT hyper) showed differential expression between tumor types. The expression of selected TFs enriched in tumor-specific DMRs are visualized with boxplots (*,**, and *** refer to P < 0.05, P < 0.01, and P < 0.001, respectively). Upper part is for RNA-seq data and lower for GSE42658 array data. **B.** Pluripotency related TFs. The expression of selected TFs enriched in tumor-specific DMRs are visualized with boxplots (*,**, and *** refer to P < 0.05, P < 0.01, and P < 0.001, respectively). Left part is for RNA-seq data and right for GSE42658 array data.

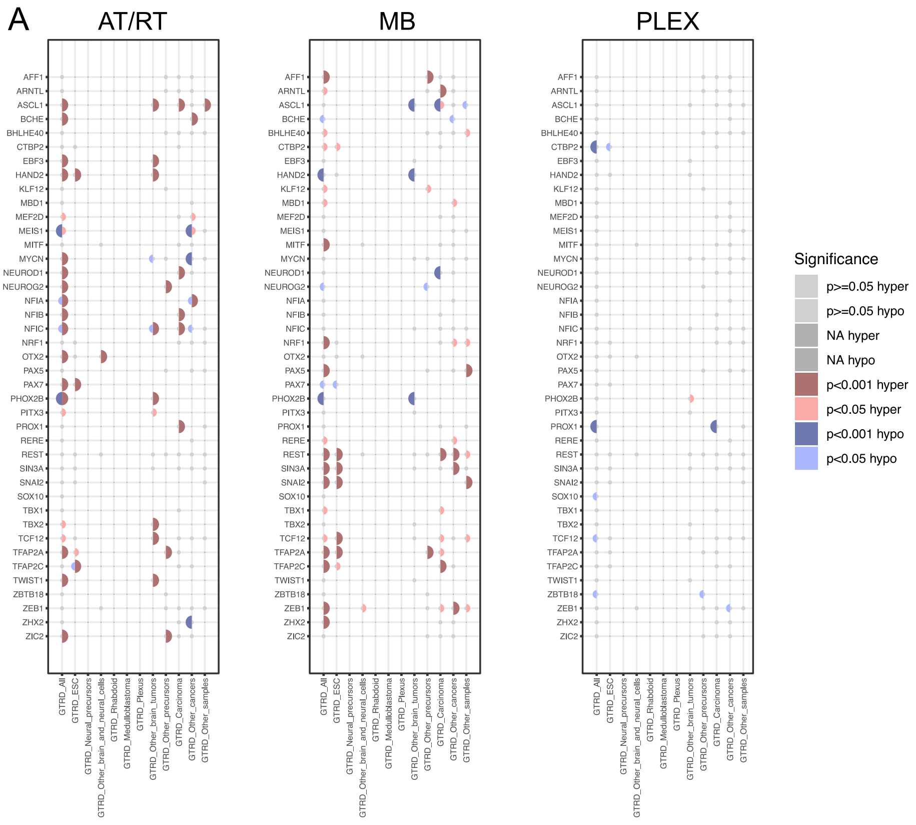


**Supplementary Figure 6.**

**A.** The enrichment of neural TF binding sites when GTRD data was split based on the sample-of-origin in the listed categories (bottom). Dot is not marked when the given TF is not measured in the given GTRD category

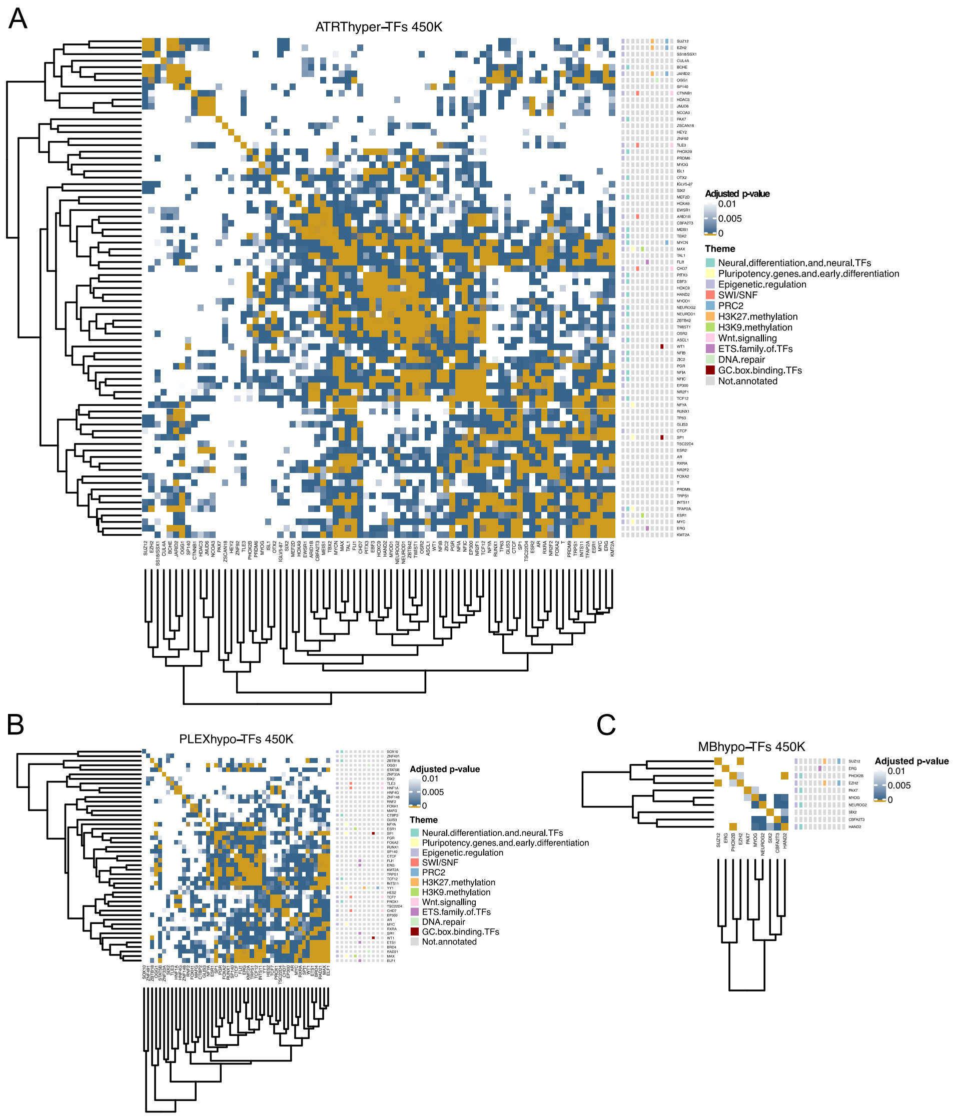


**Supplementary Figure 7**.

**A.** Colocalization heatmap of TFs with binding sites enriched to regions hypermethylated in AT/RT. All the adjusted p-values 0.001 or higher are marked in white. **B.** Same as A. but for regions hypomethylated in PLEX. **C.** Same as A. but for regions hypomethylated in MB.

A

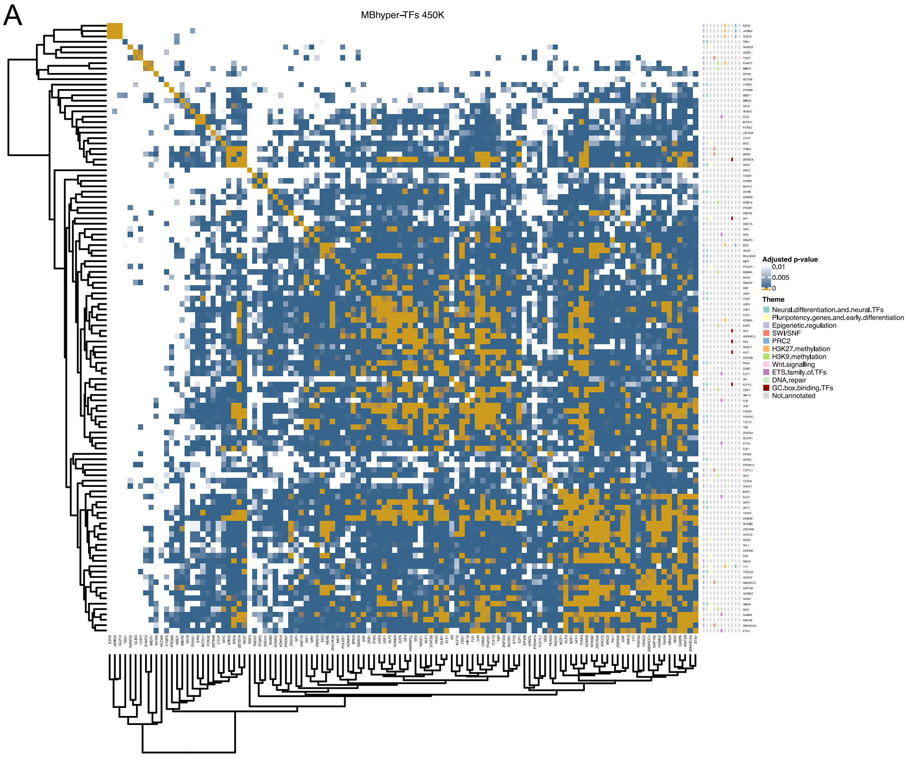


**Supplementary Figure 8.**

**A.** Colocalization heatmap of TFs with binding sites enriched to regions hypermethylated in MB. All the adjusted p-values 0.01 or higher are marked in white.

**A**

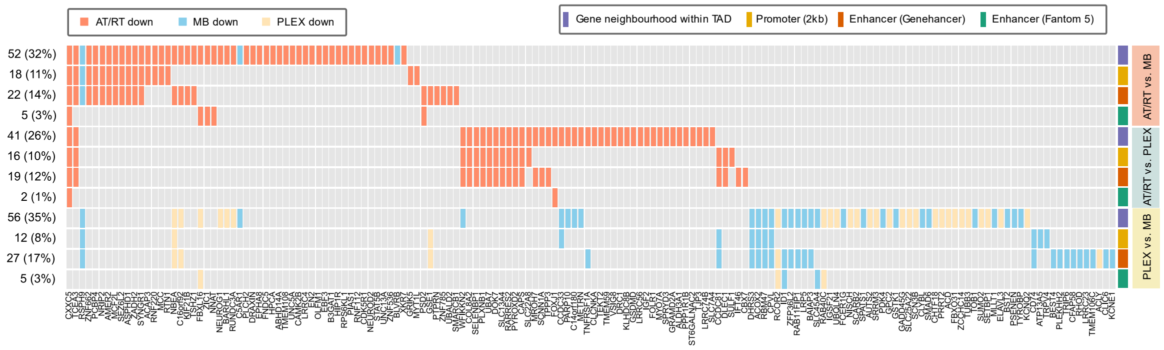


**Supplementary Figure 9.**
**A.** Oncoprint showing the DM-DE genes, which are differentially expressed in tumor comparisons, and the direction of differential gene expression. For each comparison, different possible DMR locations (Gene neighbourhood within TAD, Promoter (2kb), Enhancer (Fantom 5), Enhancer (Genehancer)) are visualised separately.

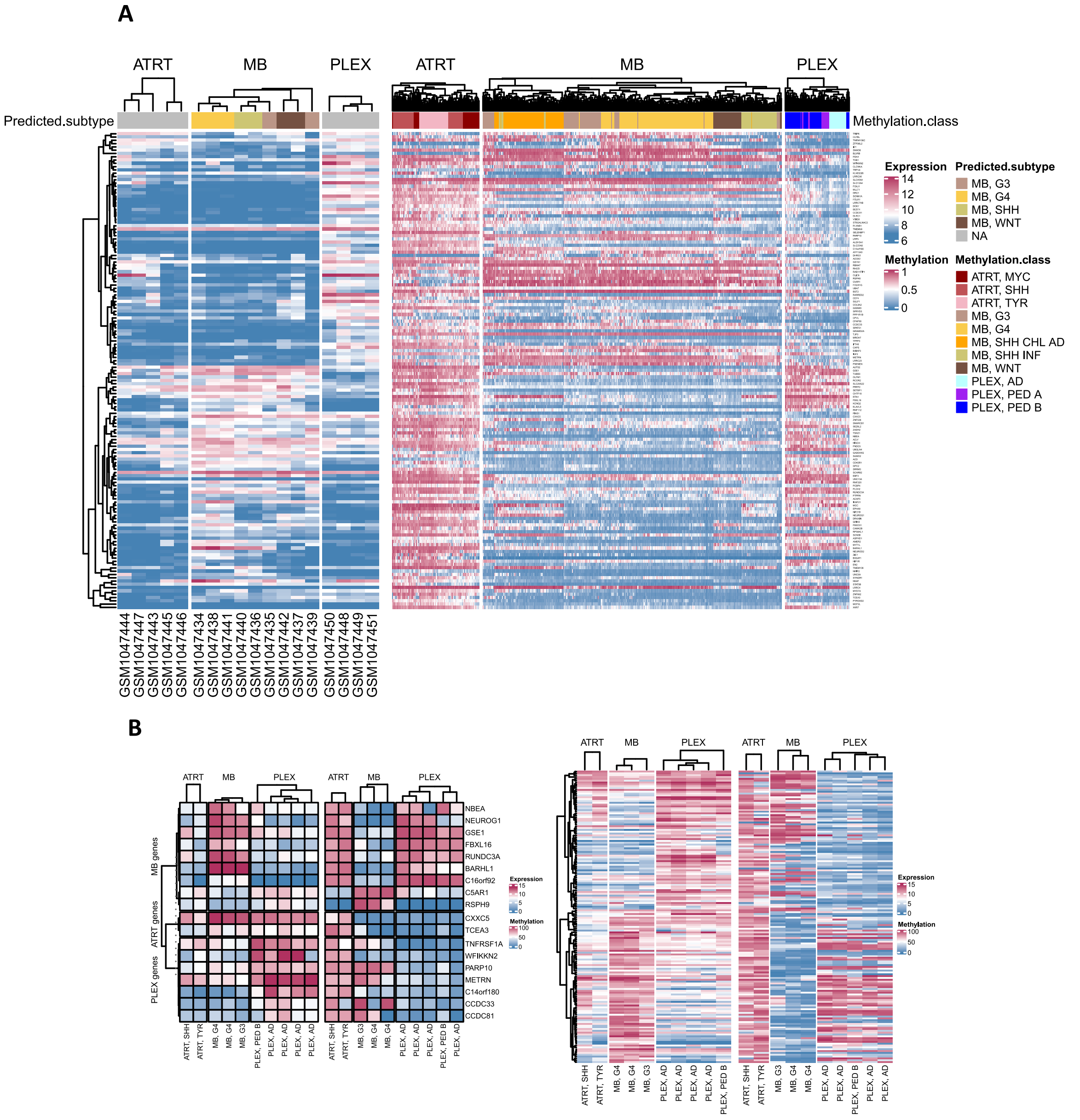


**Supplementary Figure 10.**
**A.** Expression and methylation heatmaps for DM-DE genes using public microarray data. Expression on the left and methylation on the right. **B.** Expression and methylation heatmaps for DM-DE genes using sequencing data (RNA-seq, RRBS). Expression on the left and methylation on the right.

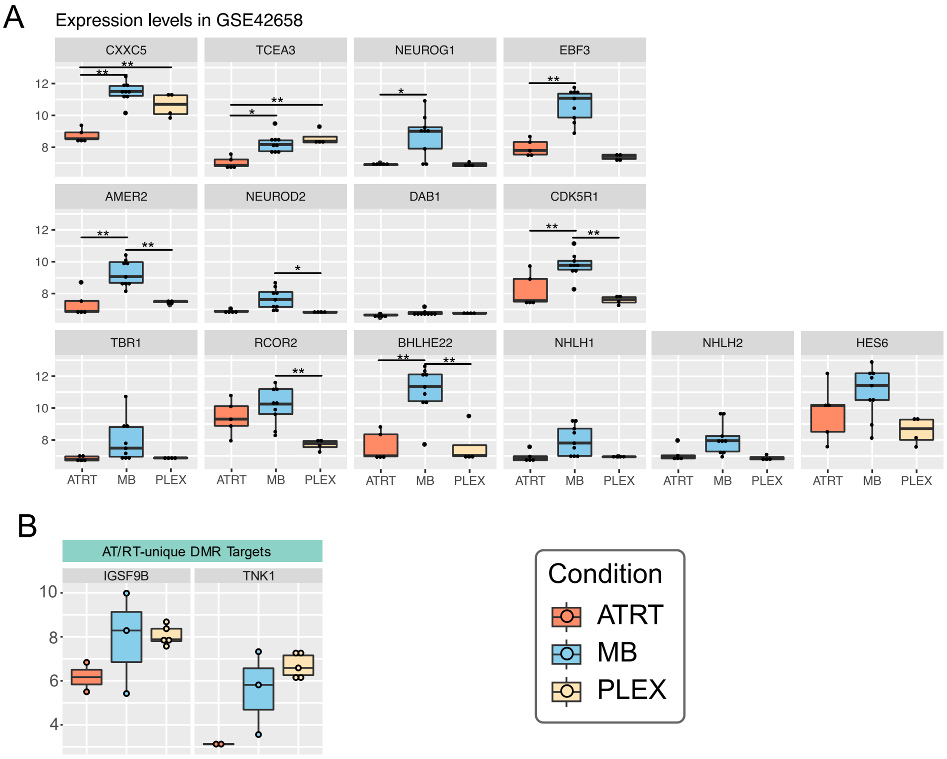


**Supplementary Figure 11. A.** The expression of genes presented in Figure 3D and NEUROG/NEUROD target genes from microarray data (GEO accession GSE42658). **B.** The expression of AT/RT-unique DMRs with EZH2 binding site target genes from RNA-seq data.

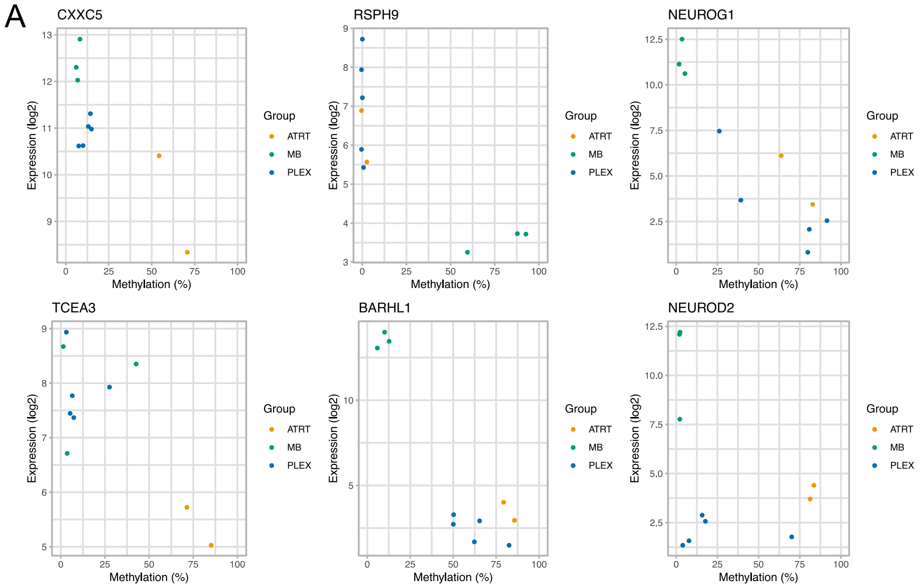


**Supplementary Figure 12. A.** Methylation-expression correlation plots for selected genes
